# Engineering antigen-specific tolerance to an artificial protein hydrogel

**DOI:** 10.1101/2023.09.23.559120

**Authors:** Peter B. Rapp, Joshua A. Baccile, Rachel P. Galimidi, Jost Vielmetter

## Abstract

Artificial protein hydrogels are an emerging class of biomaterials with numerous prospective applications in tissue engineering and regenerative medicine. These materials are likely to be immunogenic due their frequent incorporation of novel amino acid sequence domains, which often serve a functional role within the material itself. We engineered injectable “self” and “non-self” artificial protein hydrogels which were predicted to have divergent immune outcomes *in vivo* on the basis of their primary amino acid sequence. Following implantation in mouse, the non-self gels raised significantly higher anti-gel antibody titers than the corresponding self gels. Prophylactic administration of a fusion antibody targeting the non-self hydrogel epitopes to DEC-205, an endocytic receptor involved in T_reg_ induction, fully suppressed the elevated antibody titer against the non-self gels. These results suggest that the clinical immune response to artificial protein biomaterials, including those that contain highly antigenic sequence domains, can be tuned through the induction of antigen-specific tolerance.

## INTRODUCTION

Engineered proteins containing “artificial” (i.e. unnatural or *de novo* designed) amino acid sequences are playing increasingly important roles in human medicine. Prominent examples include fast-acting derivatives of human insulin,^1-2^ GLP-1R agonists, enzyme replacement therapies such as Cerezyme,^3^ and numerous engineered monoclonal antibodies such as the recent series of cancer checkpoint inhibitors, including Ipilimumab (anti-CTLA-4) and others.^4-5^ These designer proteins are routinely mutated at multiple sites in order to optimize their target affinity or bioavailability (e.g. Cerezyme H495R, Insulin “Aspart” P28D).^1-3^ Moreover, chimeric monoclonal antibodies can sometimes carry whole sequence domains derived from non-human species.^5^ The development of novel artificial protein therapeutics continues at a remarkable pace, with more than one hundred such agents already approved and in use across the US and Europe.^6-7^ Engineered protein biomaterials, including nanoparticles^8^ and hydrogels^9-14^, also lie on the clinical horizon, holding particular promise in the areas of enhanced wound healing and tissue reconstruction.^15-19^

As the clinical significance of engineered proteins continues to rise, attention must be paid to the immunological repercussions of administering these novel agents.^5, 20-22^ Adaptive immunity is a complex and multifaceted host response that is trained to generate neutralizing antibodies against foreign proteins.^23-26^ Multiple clinical studies report strong correlations between antibody emergence and a loss in therapeutic protein efficacy, an outcome which is often exacerbated by repeat administration.^27-33^ Certain neutralizing antibodies can further trigger autoimmunity by cross-reacting with endogenous host proteins.^34^ To prevent the immune-mediated rejection of artificial protein therapies, it is imperative that the emergence of neutralizing antibodies be either *a*) passively avoided or *b*) actively suppressed.

Reducing the epitope content of artificial proteins through sequence modification, a process known as deimmunization, represents a passive approach to avoiding immune activation. This strategy is predicated on the notion that antibodies are more likely to develop against highly “non-self” proteins, i.e., those with significant sequence divergence from those of the “self” host proteome. Deimmunization has several drawbacks. First, sequence modification of potential epitopes may not always be possible, as it could result in loss of function. Second, clinical data indicates that deimmunization sometimes fails to prevent host sensitization. For example, the antibody response rate to recombinant human IFN*β*-1b containing only a single C17S mutation (“Betaseron”, unglycosylated and expressed in *E. coli*) is 45%. Fully glycosylated human IFN*β*-1a with no sequence modifications (“Rebif”, expressed in Chinese Hamster Ovary “CHO” cells) has a reduced but still significant response rate of 13–24%.^30^ Attempts to humanize engineered antibodies via replacement of Fc- and V-chain fragments with human-like sequences lowers but does not completely eliminate the risk of sensitization.^23, 35-36^

An alternative strategy to passive immune avoidance is active immune suppression or tolerance. This approach aims to induce specific immune non-responsiveness to the therapeutic protein itself, thereby enhancing or rescuing its ultimate clinical efficacy. The ideal tolerization protocol would be general with respect to protein sequence and would thus be applicable to any therapeutic protein of interest. Such tolerance protocols could be particularly enabling for the field of modern tissue engineering, where the routine incorporation of artificially designed sequences renders the resultant biomaterials both highly functional as well as immunologically suspect. Recently, some studies have begun to explore the immunological repercussions of biomaterial delivery, often with the goal of harnessing biomaterials as vaccine adjuvants to augment or induce immunity.^13, 21, 37-47^ For applications such as long-term graft implantation and tissue replacement, however, tolerance is likely to be the preferred immunological outcome.^21, 48-49^

Here we report the demonstration of a broadly applicable approach for inducing antigen-specific tolerance to protein-based biomaterials. Through iterative screening and mutagenesis of a self coiled-coil domain native to mouse, we engineered an artificial non-self coiled-coil domain that was predicted to be strongly immunoreactive in mouse. We used this domain to assemble injectable protein hydrogels that resemble model non-self artificial tissues. The injected non-self hydrogels were found to raise high levels of murine anti-gel antibody titers, relative to that of a control self gel. Remarkably, fusion antibody-mediated prophylactic delivery of the non-self coiled-coil domain to DEC-205, an endocytic immune receptor known to be involved in regulatory T cell (T_reg_) induction, fully suppressed the emergence of elevated antibody levels in response to hydrogel injection. These results suggest that it should ultimately prove possible to build host tolerance to any artificial protein-based biomaterials of clinical interest, irrespective of its sequence.

## RESULTS AND DISCUSSION

### Evolution of an antigenic coiled-coil domain

We set out to develop an artificial protein sequence domain that would fulfill two criteria for the purposes of our study: 1) the domain itself would promote the efficient self-assembly of multifunctional protein biomaterials and 2) the sequence would be non-self, i.e., highly divergent from any self protein coding sequences (genomic exons) found in the mouse genome. The latter criterium would allow us to isolate the sequence determinants of immunoreactivity in the resultant biomaterials, as well as to test various adaptive immune tolerance strategies. Artificial telechelic proteins carrying α-helical coiled-coil domains at their N- and C-termini are known to spontaneously self-assemble into water-soluble polymer networks, or hydrogels.^50-51^ An appealing physical property of these networks is their mechanical reversibility or capacity to autonomously heal. This property allows them to recover their elastic modulus after subjection to high shear, thus facilitating hydrogel delivery to specific biological compartments through simple injection.

The starting point for our design was the coiled-coil forming domain of Cartilage Oligomeric Matrix Protein (COMPcc C68S C71S, Gunasekar^52^ numbering, denoted “P”), a 42mer sequence that forms pentameric coiled-coils (**Fig. 1A**).^53-54^ COMPcc is genomic exon located on chromosome 8 of *Mus musculus*; as such, it should be naturally tolerized through clonal deletion and anergy during normal thymic development in mouse. We and others have previously used the P domain to build injectable protein hydrogels that are driven to spontaneously self-assemble via P domain self-association.^55-60^ Two Cys→Ser mutations on P prevent irreversible oxidative crosslinking of the gels through cysteine-mediated disulfide formation. Since the side chains of Ser and Cys are isosteric and COMPcc is centrally tolerized, the P domain represents a model self protein that is expected to be minimally immunogenic against a mouse background

**Figure 1.**
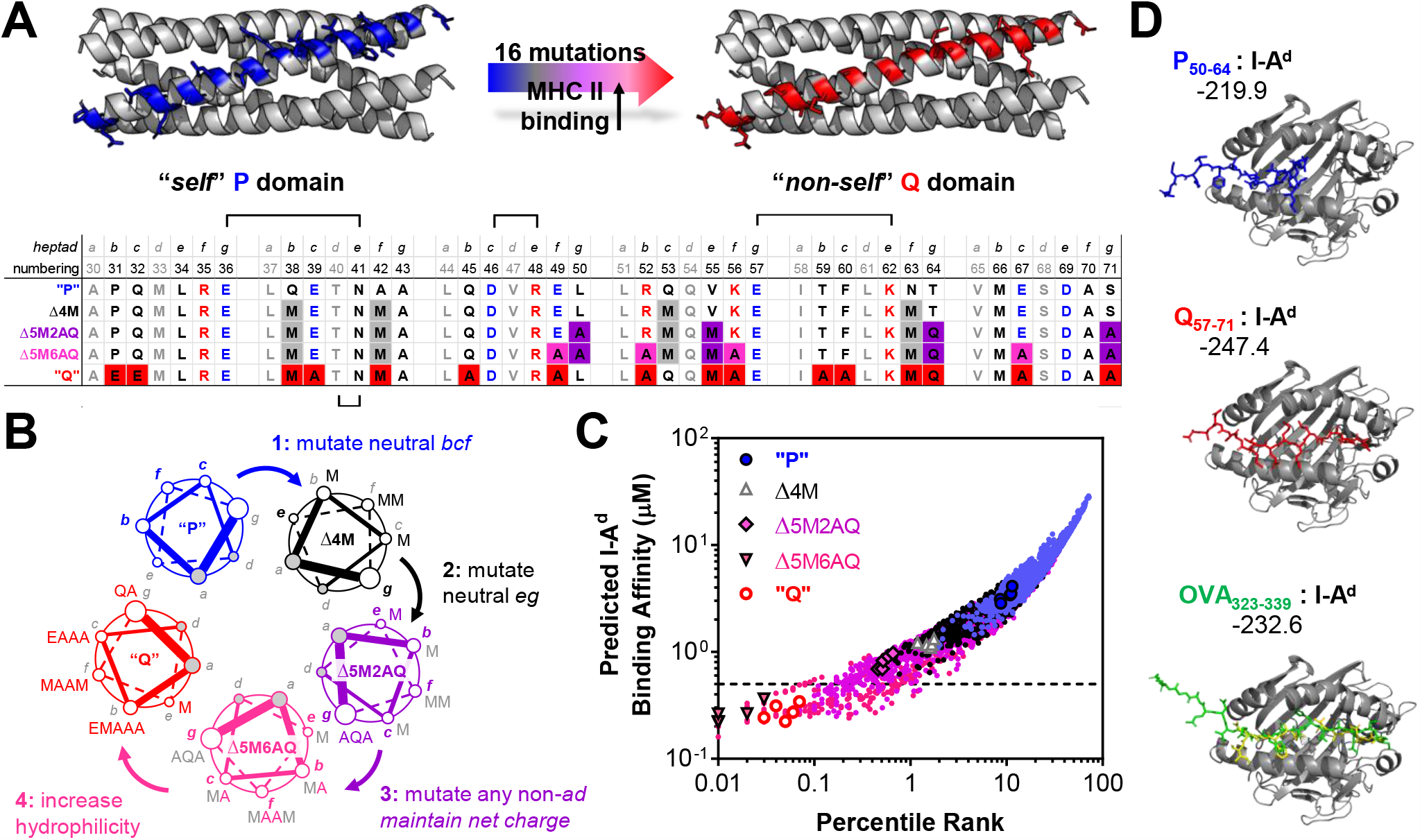
Computational design and evolution of an artificial non-self coiled-coil domain. (**A**) The P domain is a self coiled-coil derived from mouse. Mutational analysis identified a non-self Q domain carrying 16 mutations predicted to enhance the binding of proteolytic peptide fragments to Major Histocompatibility Complex II (MHC II). Sequences of the starting, intermediate and final coil sequences are shown. Lines between heptads indicate structurally important positions that were not mutated. (**B**) Overview of the mutational process (see *Main Text* for details). Each of the 4 rounds consisted of generating *in silico* sequences from all possible single point mutations (excluding Gly, Cys and Pro) at the indicated coil positions, then ranking predicted fragment immunogenicity. Mutations fixed in each round are indicated in the subsequent clockwise sector of the helical wheel diagram. (**C**) The predicted binding affinities of all unique coil-derived peptide fragments bound to MHC II haplotype I-A^d^ (*m* = 3,606) for all mutant coils screened (*n* = 412). The top 5 predictions for the coils selected and/or mutated during each round are overlayed. (**D**) Computational docking scores for the lowest-ranking (most immunogenic) Q- and P-derived peptide fragments. Q-derived peptides (red) gave better docking scores than both the P-derived peptides (blue) and the known antigenic peptide OVA_323-339_ (green). The native docking pose for the latter peptide, taken from the crystal structure (PDB 1IAO), is shown in yellow.

We performed extensive *in silico* mutational analysis on the P domain, guided by sequence comparisons to known epitopes, in order to amplify its predicted immunogenicity (**Fig. 1B** and **Fig. 1C**). The critical moment in the activation of CD4+ T cells by dendritic cells (DCs) is the display of proteolytically digested peptide fragments on a highly polymorphic cell surface receptor known as Major Histocompatibility Complex Class II (MHC II). Crystal structures of MHC II domains have revealed a set of hydrophobic “anchor” residues in the peptide binding groove that trap the processed peptide in the binding pocket.^61^ A large number of antigenic peptides have been identified in recent years, the sequences of which are publicly available on the NIH Immune Epitope Database (IEDB).^62^ The efficiency of peptide presentation on MHC II can be estimated from a positional analysis of different peptide fragments within the binding groove, and from sequence comparisons to known epitopes.

Using predictive neural-networks hosted by the IEDB, we screened 412 mutant coils in 3,606 unique peptide : MHC binding interactions, and ranked their expected immunogenicity against several hundred known database antigens (**Fig. 1C**).^63-65^ Through 4 rounds of mutation, selection, and diversification, we evolved a new coiled-coil domain designated “Q”. Relative to P, the Q domain carries 16 sequence mutations, each intended to be minimally destabilizing to coiled-coil folding. Peptidic fragments derived from Q, however, are predicted to have mid-nanomolar level binding to the MHC II receptor I-A^d.^ This receptor variant is present in BALB/c, a common inbred mouse strain. As an independent predictive test for the increased immunogenicity of Q, we performed blind peptide-protein docking to the MHC II cleft using HPEPDOCK.^66^ The most immunogenic IEDB-predicted peptide epitopes derived from P and Q were permitted to globally sample the empty I-A^d^ structure in a flexible and unbiased fashion. Consistent with the design criterion, Q-derived peptides had significantly higher docking scores (i.e., lower relative docking energies) than the P-derived peptides (**Fig. 1D** and **Table S1**). HPEPDOCK also correctly docked the known peptide antigen OVA_323-339_ to I-A^d^ with a docking score similar to that of Q_57-71_, further supporting the validity of our computational antigen design approach. Intriguingly, the Q domain bears no homology to any known protein sequence as evidenced by a null search result in BLASTP.^67^ Strong predicted interactions with I-A^d^ are a unique property of Q, as other helical multimerization domains, including viral-derived coiled-coil domains such as from influenza A/B, have weak predicted interactions with this MHC II haplotype (**Table S2**).

### Physical characterization of artificial protein hydrogels

To test whether the mutant Q domain was competent to drive biomaterial self-assembly, we cloned and expressed a recombinant “QXQ” block copolymer. This telechelic protein comprises two associative Q domains flanking a flexible 144mer sequence taken from the engineered XTEN protein, here denoted as “X” (**Fig. 2A**). Similar in concept to *poly*-ethylene glycol (PEG), XTENs were originally developed by Stemmer et al. as a genetically encodable tool for half-life extension of recombinant biologics.^68^ They have been demonstrated to be immunologically inert due to the absence of hydrophobic anchor residues or protease cleavage sites in their primary amino acid sequence. As such, any immune responses observed against a QXQ hydrogel architecture may be presumed to be directed principally against the antigenic Q domain. We also prepared the corresponding “PXP” block copolymer using the unmodified P domain (see **Table S3** and **Table S4** for expressed protein and plasmid sequences, respectively).

**Figure 2.**
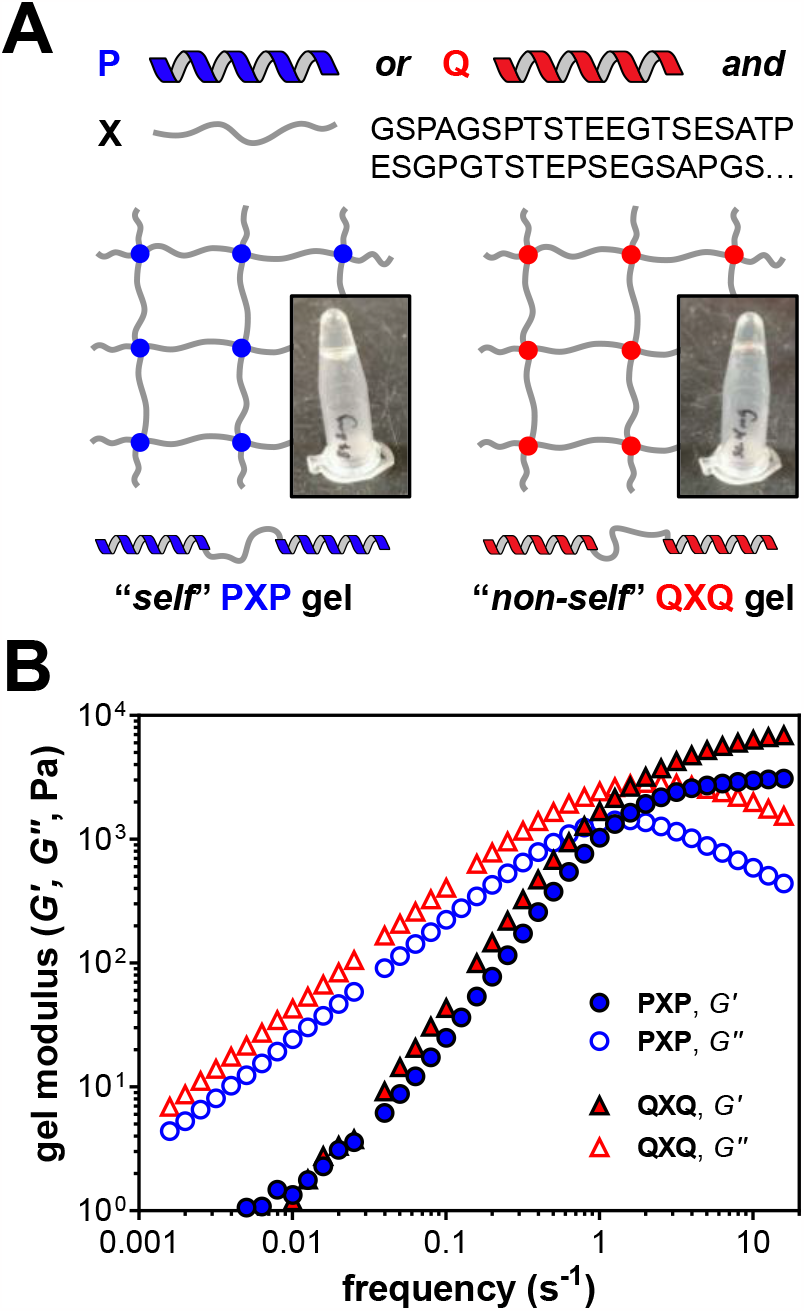
Dynamic material properties of the self and non-self gels. (**A**) The PXP and QXQ proteins spontaneously formed transparent hydrogels upon addition of phosphate buffer to the lyophilized protein. Shown are 10% (*w*/*v*) gels suspended in 100 mM P_i_ pH 7.4. (**B**) Oscillatory shear rheology of both gels revealed matching reversible viscoelasticity. Frequency sweeps were performed at a fixed strain amplitude of 1% between 0.01 and 100 rad s^-1^. The gels were incubated at a fixed temperature of *T* = 37 °C

Both the PXP and QXQ proteins spontaneously formed optically clear, reversible hydrogels upon addition of neutral phosphate buffer to the lyophilized protein (**Fig. 2A**). Qualitatively, the physical behavior of these gels is similar to P(E_*n*_P)_*m*_-type multiblock gels carrying an elastin-like “E” polypeptide midblock.^55, 57, 59-60, 69^ In contrast to elastin-like gels, however, the XTEN-based gels do not display an inverse thermal phase transition temperature (LCST) upon heating. To quantitively compare the PXP and QXQ gels, their dynamic mechanical moduli were assessed through oscillatory shear rheology (**Fig. 2B**). Both materials exhibited classical Maxwellian viscoelastic reversibility, with an elastic modulus (*G*’) dominating at high frequency and a viscous modulus (*G*’’) dominating at lower rates of material deformation. At 37 °C, the terminal modulus of both materials is similar, and lies between *G*_∞_ = 3.2 – 4.7 x 10^3^ Pa. The characteristic relaxation rates are also comparable, with ω_c_ = 1.9 – 5.4 s^-1^ at *T* = 37 °C (**Fig. S1**). In further support of multimerization competency of the Q domain, dilute solutions of QXQ displayed increased helicity relative to PXP, as evidenced by circular dichroism spectroscopy (QXQ: θ_222_/θ_208_ = 0.79; PXP: θ_222_/θ_208_ = 0.66) (**Fig. S2**).

In the process of developing the Q-based XTEN gels, we screened several additional P-derived mutant coiled-coils for relevant mechanical properties. The additional coils were cloned and expressed as coil-elastin-coil (CEC) fusion constructs, with the mutant coils flanking an elastin-like midblock (see *Supporting Information* **Fig. S3, Table S5**, and **Table S6** for additional details). These CEC-type materials exhibited a wide range of terminal stiffnesses, relaxation rates, and phase separation temperatures. Notably, whereas PEP gels had a LCST of >95 °C, QEQ gels exhibited coacervation at a much lower temperature of 45 °C. Moreover, some coils with high predicted immunogenicity (e.g., 5M6AQ, Coil #6), exhibited phase separation at or below room temperature when cloned into the elastin framework. Increased predicted immunogenicity of the mutant coils was generally correlated with increasing mutational distance from P, and inversely correlated with phase separation temperature.

### Immunological response to hydrogel implantation

Gels assembled from the PXP and QXQ proteins represent model self and non-self biomaterials, respectively. PXP is comprised of self protein and the nonimmunogenic XTEN domain, whereas QXQ carries a highly antigenic non-self domain engineered specifically to provoke a strong adaptive immune response. As such, PXP and QXQ gels arguably represent “best-case” and “worst-case” immunological scenarios, respectively, for clinical grade engineered protein biomaterial implants. We proceeded to probe the humoral adaptive immune response to these two hydrogels using a novel injectable hydrogel challenge model (**Fig. 3A**). Both gels are shear thinning and could be readily injected into the mouse subcutaneous cavity, whereupon they immediately reassembled to form a cohesive protein depot (**Fig. 3B**). Because the mechanical properties of the two gels are similar (*cf*. **Fig. 2**), differences in immunological outcomes following implantation may be reasonably attributed to sequence differences between P and Q.

**Figure 3.**
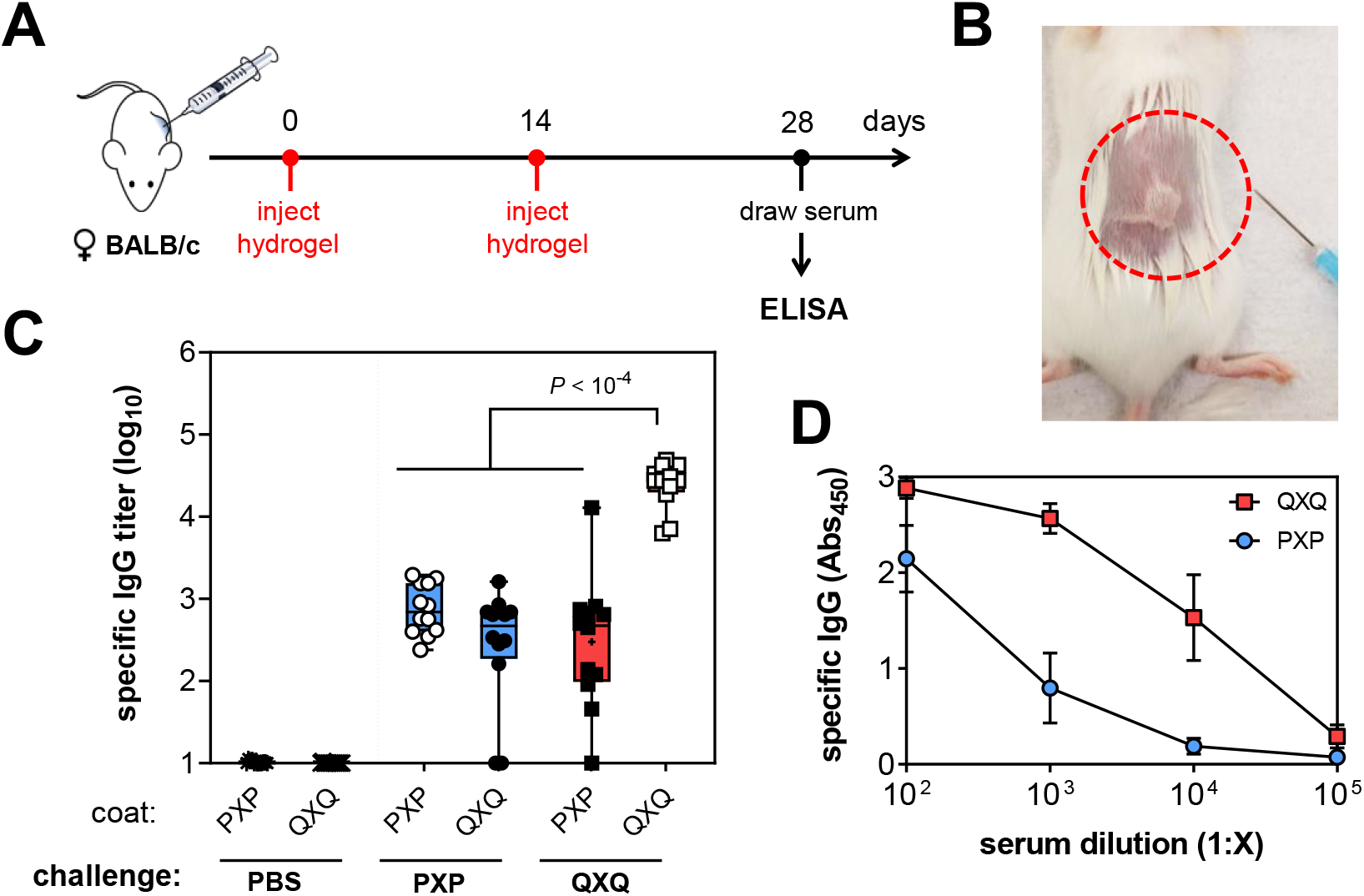
Sequence-dependent immunoreactivity of injected artificial hydrogels. (**A**) Self (PXP) and non-self (QXQ) hydrogels were implanted into mice using a novel injectable hydrogel challenge model. Female BALB/c mice (8-10 weeks old, *n =* 4 mice/group) were challenged on d0 with subcutaneous hydrogel implantation (10% *w*/*v*, 5 mg in 50 μL of 100 mM P_i_ pH 7.4), then boosted on d14 with a second gel injection. Serum was recalled for ELISA on d28. (**B**) Photograph of the hydrogel implantation site immediately following injection. Mechanical reversibility permits the gel to stay assembled after being extruded from a 22G needle. (**C**) Specific IgG titers against PXP and QXQ were determined by indirect ELISA. Sera from mice injected with the indicated challenge proteins were incubated in wells coated with either PXP or QXQ. No signal was seen in wells containing no adsorbed antigen (uncoated). Mice challenged with QXQ had 30-fold higher ELISA signal against QXQ than did PXP-challenged mice against PXP. Shown are all sera results from three independent biological replicates of 4 mice / group (3 x *n* = 12 total mice). (**D**) Dilution series of sera from PXP- and QXQ-challenged mice reacted against the same coat protein.

The BALB/c mouse strain carries the MHC II haplotype I-A^d^, and the high epitope content of Q is specific to this receptor. BALB/c was therefore selected as the appropriate mouse model for our challenge studies. We hypothesized that within this strain, protein biomaterials embedded with Q domains would be highly immunogenic, such that immune phenotyping and tolerance induction protocols could be benchmarked. C57BL/6, another common inbred mouse strain, carries the related haplotype I-A^b^. However, neither P nor Q was predicted by IEDB analysis to interact with the I-A^b^ haplotype. C57BL/6 therefore serves as a reasonable control strain for the sequence-dependence of the immune response to P *vs*. Q.

Female BALB/c mice were challenged twice with hydrogel injections on d0 (prime) and d14 (boost), followed by sera recall and analysis by indirect ELISA (**Fig. 3A** and **3B**). Importantly, this assay queries the *antigen specificity* of the subsequent immune response as opposed to the total antibody content of the serum. Whereas sera from PXP-challenged mice exhibited intermediate IgG titers against the challenge protein, sera from QXQ-challenged mice exhibited significantly stronger antibody titers (**Fig. 3C** and **Fig. 3D**). In particular, ELISA signal from QXQ-coated wells incubated with QXQ challenge sera was 30-fold higher than the corresponding PXP-coated wells incubated with PXP challenge sera, representing a specific IgG titer difference of 1.5. The high titer against QXQ was similar in magnitude to that seen when mice were challenged with Ovalbumin admixed to Alum, a immunogen known to be potent in BALB/c (**Fig. S4**).

To further probe the specificity of the humoral IgG response, we cross-reacted challenge sera with the opposite coat protein (**Fig. 3C**). This analysis revealed that the elevated titer was specific to epitopes located in the Q domain as opposed XTEN, since PXP-coated wells incubated with QXQ challenge sera returned intermediate titer levels comparable to that seen with PXP challenge sera alone. Moreover, this difference in titers was strain-dependent and specific to the BALB/c mouse. C57BL/6 mice challenged with either PXP or QXQ produced similar titers (**Fig. S5**). Taken together, these *in vivo* results establish that non-self QXQ hydrogels induce an elevated level of sequence-dependent immunoreactivity relative to that of the self PXP hydrogels. These results are also consistent with the computational design criterion for the Q domain as a *de novo* T-cell-dependent biomaterial antigen. Although PXP does not contain sequence fragments that were predicted to be strongly immunogenic, we consistently observed low-to-intermediate IgG titers against this material (e.g. **Fig. 3C**). One possible driver of this phenotypic response is endotoxin or lipopolysaccharide (LPS), a ubiquitous contaminant of *E. coli*-derived recombinant proteins. Endotoxin can trigger potent immune responses through engagement of Toll-like receptor 4 (TLR-4). To exclude the possibility that the differences between PXP and QXQ were dependent on differential endotoxin levels within these two protein preparations, we cloned a stable *CHO*-based mammalian cell line for endotoxin-free expression of QXQ (**Fig. S6**). QXQ protein expressed from *CHO* had nearly undetectable levels of endotoxin (<0.1 EU/mg) as compared to *E. coli* expressed QXQ (<10 EU/mg). However, despite the significant differences in endotoxin levels, anti-QXQ titers against the *CHO*-derived proteins were equivalent to those against *E. coli*-derived protein in the hydrogel challenge model (**Fig. S6**).

Alternative explanations for the intermediate PXP titers are 1) weak (and/or T-cell-independent) underlying immunoreactivity of some part of the PXP construct (e.g. 6xHis), or 2) the presence of low levels of contaminating antigenic background proteins from the *E. coli* proteome in each recombinant batch. Consistent with the first hypothesis, injection of XTEN protein alone (without Q but carrying 6xHis at both N- and C-termini) also induced intermediate IgG titers, and sera from both PXP- and QXQ-challenged mice cross-reacted with *E. coli*-derived 6xHis-tagged GFP coat protein (**Fig. S7**). Consistent with the second hypothesis, the cross-reactivity against *E. coli*-derived GFP coat protein was undetectable when challenge sera from *CHO*-derived QXQ was plated (**Fig. S7**).

### Tolerance induction to an artificial protein hydrogel

The observation of sequence-dependent immunoreactivity against an artificial protein hydrogel raises the question of whether it is possible to attenuate this “anti-gel” antibody response in a general but sequence-specific manner. Although broadband (i.e., antigen-nonspecific) immunosuppressants such as cyclosporine, methotrexate and rapamycin are widely used to prevent the rejection of organ transplants,^70^ such chemotherapeutics carry the serious unwanted side-effect of compromising the host immune response to harmful pathogens and increasing the risk of infection. Antigen-specific tolerance, by contrast, would deactivate the immune system towards only a subset of functional antigens, thereby enabling the implantation or grafting of any artificial protein biomaterial irrespective of its sequence. This clinical possibility represents a shared landmark goal for the fields of immunoengineering and biomaterials medicine.^71^

Reverse vaccination represents one strategy by which antigen-specific tolerance to artificial implants could be induced. In this strategy, a dose of antigen is prophylactically administered to the host in such a manner that the antigen is immunologically sanctioned as self, such that future challenges with the same antigen do not provoke a subsequent inflammatory response. Tolerance induction has been explored in a wide variety of immunologic contexts, often with the goal of preventing or reversing autoimmunity to critical host proteins, e.g. β-cell-derived antigens in the case of type 1 diabetes (T1D).^72^ Broadly speaking, reverse vaccination aims to achieve either 1) anergy/deletion of specific auto-reactive effector T cell (T_eff_) and B cell (B_eff_) clones, or 2) induction of antigen-specific regulatory T cells (T_reg_) that actively suppress effector function against specific immunoreactive epitopes. Although the detailed biomolecular mechanisms of T_reg_-mediated immunosuppression are diverse and context-dependent, induction of T_regs_ is associated with improved immunological outcomes in autoimmunity, allergy, organ transplantation, and the maintenance of feto-maternal tolerance.^73-79^

We surveyed antigen-specific tolerance strategies^72, 80-84^ with the goal of identifying a tolerance induction protocol that would attenuate the anti-gel antibody response seen against non-self biomaterials such as QXQ, conceivably through the *in vivo* induction of antigen-specific T_regs_. Carbodiimide-mediated crosslinking of peptides to the surface of splenic leukocytes has been shown to induce systemic tolerance through the induction of T_regs_, but requires the use of autologous (i.e., host-derived) cells.^85^ Oral feeding of antigen^86-90^ as well as certain commensal bacteria^91^ are known to promote T_reg_ development for the maintenance of intestinal health, but the gut compartmentalization of these natural tolerance mechanisms render them difficult to co-opt for the generation of systemic tolerance. Other noteworthy tolerance phenotypes have been induced via nanoparticles loaded with antigen or decorated with peptide-MHC,^92-94^ coupling of antigens to the surface of erythrocytes,^95-96^ as well as activation of inhibitory Siglec receptors (e.g. CD22) by antigens decorated with sialic acid ligands.^97-101^ Recently, therapeutic tolerance via so-called “inverse vaccination” using antigens chemically conjugated to polymeric *N*-acetylgalactosamine (pGal) has also been explored.^102^

One conceptually straightforward approach to tolerance induction is to perform natural antigen presentation in the absence of ongoing inflammation or costimulatory ligands. Under these conditions, the default immunologic response to exogenous antigen is adaptive tolerance.^103^ Nussenzweig and Steinman demonstrated the importance of immature dendritic cells (DCs) for peripheral tolerance induction in the immunological steady state.^104-106^ These studies identified DEC-205, a C-type lectin domain enriched on certain immature DCs, as an endocytic gatekeeper receptor for steady state tolerance induction.^107-110^ Antigens delivered to DEC-205 are targeted to the late endosome and undergo efficient antigen display on MHC II, ultimately leading to robust T-cell tolerance via antigen-specific T_reg_ induction.^111-115^ We hypothesized that DEC-205-mediated endocytosis could be harnessed to induce systemic tolerance to artificial protein hydrogels via reverse vaccination (**Fig. 4A**).

**Figure 4.**
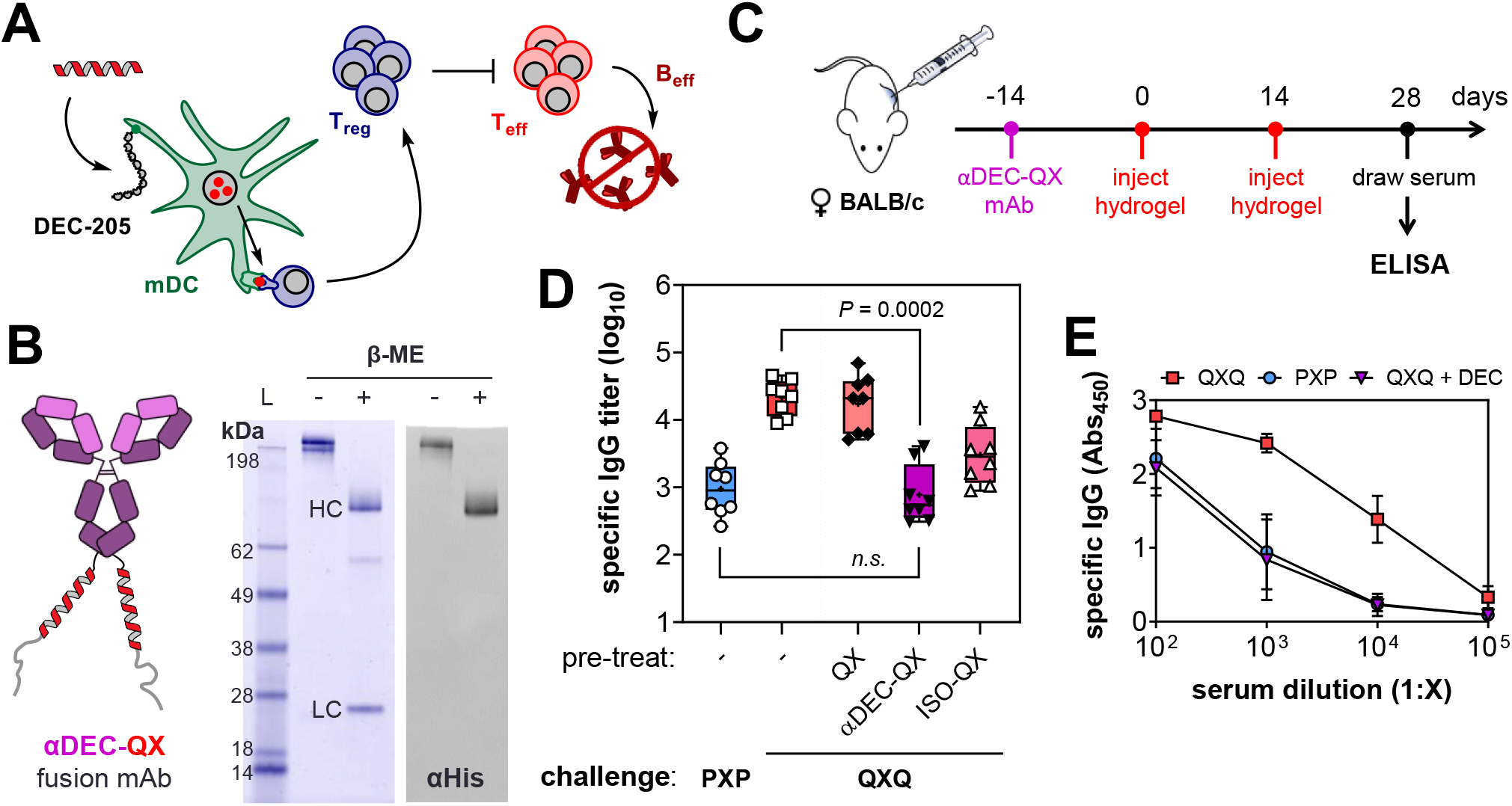
Induction of immune tolerance to QXQ hydrogels through steady state targeting of DEC-205. (**A**) Antigens uptaken by DEC-205 undergo deep endosomal penetration and high-efficiency display on MHC II. In the immunological steady-state (i.e., absence of costimulation), this induces antigen-specific regulatory T cells (T_reg_) that attenuate the antibody response (B_eff_) through active suppression of effector T function (T_eff_). (**B**) Q and XTEN domains were fused in series to the HC of αDEC-targeting antibody clone NLDC-145 to prepare αDEC-QX (tagged with C-terminal 6xHis). (**C**) Female BALB/c mice (8-10 weeks old, *n =* 4/group) were prophylactically administered αDEC-QX conjugate (10 μg, IP) or an isotype control conjugate once on d-14, followed by SC hydrogel injections on d0 and d14. Sera was recalled on d28 and assayed by ELISA. (**D**) Specific IgG titers against PXP and QXQ were determined by indirect ELISA. Mice receiving prophylactic αDEC-QX had significantly reduced specific antibody titers against non-self QXQ following the hydrogel challenge. These titers were suppressed to the levels seen against the self PXP gel. Shown are all sera results from two of four independent biological replicates at 4 mice / group (2 x *n* = 8 mice / group representative of 16 total mice). (**E**) Dilution series of sera from PXP and QXQ-challenged mice reacted against the respective coat protein, compared to QXQ-challenged mice tolerized with αDEC-QX.

To test this hypothesis, we obtained cDNA for the DEC-205-specific antibody clone NLDC-145, which has previously been shown to deliver model antigens to DEC-205.^111^ We then cloned a novel fusion antibody by placing the “Q” and “XTEN” domains at the C-terminus of the NLDC-145 antibody heavy chain (HC). The resultant “αDEC-QX” antibody carries the putative T-cell determinants of QXQ immunoreactivity, as well the 6xHis epitope, within its primary amino acid sequence (see **Table S3** and **Table S4** for expressed protein and plasmid sequences, respectively). The fusion construct was expressed and purified from HEK-293-6E cultures, and assembly of the full-length (HC and LC) antibody was confirmed by anti-His blotting (**Fig. 4B**). As a targeting control, we also prepared an isotype-matched (IgG1) fusion “ISO-QX” with no affinity for DEC-205.

Next, female BALB/c mice were prophylactically administered 10 μg of either αDEC-QX, ISO-QX, or the nonconjugated QX domain, followed by immune challenge with QXQ hydrogel injected on d0 (prime) and d14 (boost). As before, sera were recalled on d28 for analysis of antigen-specific titers by indirect ELISA. Remarkably, mice which received the αDEC-QX fusion prior to implantation of QXQ gels were resistant to the development of elevated titers against QXQ (**Fig. 4D** and **Fig. 4E**). Moreover, the titer level against QXQ in these mice was indistinguishable from that of the titer level against PXP in mice that had been challenged with PXP gels. Administration of QX domain alone had no effect on the resultant level of antibody, whereas the ISO-QX control partially attenuated the high titer response to QXQ. This is consistent with other observations that long-circulating antigen (e.g., as carried by albumin) can be weakly tolerogenic.^116^ Isotyping experiments further confirmed that antigen-specific titers of IgG1, the dominant antibody isotype raised against QXQ in the injection challenge, were strongly suppressed by DEC-205 targeting (**Fig. S8**). Taken together, these results demonstrate successful induction of a striking tolerance phenotype to QXQ hydrogels through DEC-205-mediated reverse vaccination.

As already noted, the DEC-205 pathway has been previously shown to induce T_reg_-mediated tolerance to select model antigens (e.g. Ovalbumin). Our study expands the scope of DEC-205-mediated tolerance to artificial protein hydrogels, an important class of engineered biomaterials. Notably, previous tolerance studies involving DEC-205 have primarily reported cellular tolerance phenotypes and overlooked the antibody response, whereas the tolerance phenotype we describe is humoral. Intriguingly, an intermediate antibody titer remains against QXQ gels even after αDEC-QX treatment (**Fig. 4D**). This could indicate incomplete T cell tolerance generation; alternatively, T-independent immunoreactivity or trace background proteins from the recombinant expression organism may be contributing to the observed assay signal. Finally, despite the clear humoral tolerance phenotype, our evidence for T_reg_ involvement remains indirect. Additional studies are needed in order to confirm the direct involvement of T_regs_ in the present challenge model (e.g. FACS-based sorting and *ex vivo* co-culture of recalled CD25+ FoxP3+ cells).

## CONCLUSION

The ability to attenuate the immune response to specific protein antigens of arbitrary sequence (some of which are likely to possess high epitope content) has the potential to significantly broaden the design space, as well as the range of clinical applications, available to artificial protein-based biomaterials. We have shown that sequence characteristics play a critical role in the humoral response against injectable protein hydrogels. Through careful deployment of a suite of modern computational tools, we designed self and non-self protein hydrogels which differed greatly in their putative epitope content and resultant immunogenicity, despite having very similar biophysical properties. The reality of sequence-dependent immunoreactivity need not deter functional biomaterials design, however. We have further shown that antibody responses to the self and non-self gels can be normalized through prophylactic tolerance induction. This latter result conceivably opens the door to the clinical deployment of any protein-based biomaterial, irrespective of its constituent amino acid sequences. Overall, our results invite a closer inspection of the interaction between protein biomaterials and the adaptive immune system, particularly as it pertains to the functional consequences of various tolerance phenotypes.

## ASSOCIATED CONTENT

### Supporting Information

The Supporting Information is available free of charge on the ACS Publications website. Contains additional experimental details, protein sequence and expression information, protein biophysical characterization, as well as computational and assay data. This material is available free of charge via the Internet at http://pubs.acs.org.

## AUTHOR INFORMATION

### Author Contributions

P.B.R. conceived and designed the study. P.B.R., J.A.B., R.P.G., and J.V. performed the experiments. P.B.R. and J.A.B. analyzed the data and wrote the manuscript. All authors have given approval to the final version of the manuscript.

### Notes

The authors declare no competing financial interests.

## ACKNOWLEDGMENT

We wish to sincerely thank Provost David A. Tirrell (Caltech Chemistry and Chemical Engineering) and Professor Sarkis K. Mazmanian (Caltech Biology and Biological Engineering) for providing scientific mentorship and generous institutional support throughout the execution of this project. We thank the Protein Expression Center of the Caltech Beckman Institute for mammalian cell line development and protein expression.

## ABBREVIATIONS

MHC: major histocompatibility complex
IEDB: Immune Epitope Database
DC: dendritic cell
T_eff_: effector T cell
T_reg_: regulatory T cell
B_eff_: effector B cell
LPS: lipopolysaccharide
TLR-4: toll-like receptor 4

