## Supporting Information for "Engineering antigen-specific tolerance to an artificial protein hydrogel"

[1]*Division of Chemistry and Chemical Engineering,*
California Institute of Technology,
1200 E. California Blvd., Pasadena, CA, 91125

[2]*Division of Biology and Biological Engineering,*
California Institute of Technology,
1200 E. California Blvd., Pasadena, CA, 91125

*

### Table of Contents

**Materials and Methods** ... 3
*MHC docking simulations of putative antigens derived from P and Q* ... 3
*Cloning of XTEN material library* ... 3
*Expression and purification of XTEN proteins in E. coli* ... 4
*Removal of endotoxin and protein sterilization* ... 4
*Preparation of stable CHO lines* ... 5
*Expression of "endotoxin free" XTEN proteins from CHO cells* ... 6
*Cloning, expression and purification of αDEC-205-QX fusion antibody* ... 6
*Quantification of endotoxin levels* ... 7
*Oscillatory shear rheology of hydrogels* ... 8
*Immune challenge and tolerance induction protocol* ... 8
*Analysis of antibody titers by enzyme-linked immunosorbent assay (ELISA)* ... 9
*Circular dichroism spectroscopy* ... 10
**Table S1. HPEPDOCK results for top 10 P and Q antigens bound in H2-I-A$^{d}$** ... 11
**Table S2. Comparative immunogenicity analysis of other coiled-coil domains** ... 12
**Table S3. Amino acid sequences of artificial proteins, hydrogels, and antibody fusions** ... 13

### Materials and Methods

***MHC docking simulations of putative antigens derived from P and Q.*** Docking of the 10 lowest ranking (most antigenic) putative immunogens derived from P and Q (as predicted by the IEDB, cf. **Table S1**) were subject to unbiased global docking to the MHC Class II structure H2-I-A$^d$. The native peptide 17mer in PDB structure 1IAO (OVA$_{323-339}$, sequence ISQAVHAAHAEINEAGR) was manually removed from the peptide binding cleft using PyMOL. The cleaned (empty cleft) structure was then uploaded to the HPEPDOCK2.0 server. Each putative immunogenic peptide 12mer was then permitted to globally dock to any part of the structure using the default docking parameters. Following docking, the top 10 docking poses for each of the 10 peptides (ranked according to docking energy score) were qualitatively inspected for "correct" docking, i.e. the peptide was found inserted into the standard binding groove versus docked to an irrelevant site. Peptides in groove were labeled "bound" and peptides out-of-groove were labeled "unbound" (**Table S1**). The average bound docking score for all peptides (derived from either P or Q) was lower than that of the corresponding average for all unbound docking scores. Moreover, the average docking score corresponding to the top ranked model for each Q peptide was significantly lower than for the corresponding P peptide models. To validate the docking model, the OVA$_{323-339}$ peptide was independently docked in the empty cleft structure by HEPDOCK2.0. The docked peptide had a low docking score, and aligned with the corresponding "true" docking pose from the crystal structure with an RMSD of 1.70 Å$^2$ (corresponding to 11 of 17 possible residues aligned within the cleft).

***Cloning of XTEN material library.*** Construction of *E. coli* plasmids encoding for the various XTEN-based hydrogel proteins began with a modified pQE-80L-Δ*XhoI* ≡ pQE (Qiagen, USA) vector wherein the native *XhoI* site upstream of the native MCS was destroyed by site-directed mutagenesis. Genes encoding the P block (or its various mutants) and the XTEN sequence "X" were synthesized by DNA2.0 (Newark, CA), with the XTEN gene codon-optimized for maximal expression in *E. coli*. All mutant P blocks were synthesized as "gBlock" fragments to facilitate rapid cloning. Each designed gene had restriction enzyme digestion sites in the sequence 5'-*BamHI-SalI-*<u>*CDS*</u>*-XhoI-HindIII-3'* to allow for iterative insertion of multiple gene fragments. The full-length genes for each XTEN protein were prepared by recursive site-directed ligation.[1] First, a single P domain (or another mutant coiled-coil) and X were installed on different pQE vectors via digestion with *BamHI* + *HindIII* to generate pQE-P and pQE-X. An additional P domain was then inserted at the 3' end of pQE-X by digestion of the vector (pQE-X) and insert (P) with *XhoI* + *HindIII* and *SalI* + *HindIII*, respectively. Subsequent ligation generated pQE-XP with simultaneous destruction of the internal *XhoI* site, leaving the terminal 5' *SalI* (from the vector) and 3' *XhoI* sites (from the insert) intact. Repeating this procedure with pQE-P as vector and XP as insert yielded pQE-PXP. All additional XTEN genes (containing mutated P domains) were prepared in an analogous manner, starting from pQE-X and the corresponding gBlock. **Table S3** and **Table S4** contain the complete amino acid sequences of each protein prepared in this manner, as well as the corresponding coding sequences of each of the plasmids prepared. Additional mutant coils cloned in elastin "E" midblocks are listed in **Table S5** and **Table S6**.

***Expression and purification of XTEN proteins in E. coli.*** Plasmids coding for each of the XTEN hydrogel proteins were transformed into BL21 chemically competent *E. coli* (NEB, Δ*fhuA2* resistant to phage T1). Overnight cultures of transformed cells were used to inoculate 1 L flasks containing Terrific Broth (TB) (inoculation ratios were typically 1:20-50) supplemented with 100 μg ml$^{-1}$ ampicillin. Cells were grown to an $OD_{600}$ of 0.7 – 1.0 then induced with 1 mM of isopropyl β-D-1 thiogalactopyranoside (IPTG). After 4 – 6 h, bacterial cultures were harvested by centrifugation for 6 min at 10,000*g*. Cells were immediately resuspended in 8 M urea pH 8 supplemented with 100 mM phosphate, 10 mM Tris and 10 mM imidazole. Lysates were taken through two freeze-thaw cycles before being subject to high-power tip sonication (tip diameter ~ 1 cm). For sonication, 50 mL of lysate from a 1 L culture was treated at 50% maximum amplitude for 10 min in 1 sec pulse intervals (5 min total sonication time). Homogenized lysate was clarified by high-speed centrifugation (50,000*g* for 1 h) and then subjected to standard His-tag purification over Ni-NTA agarose beads (Qiagen, USA) under denaturing conditions. Following elution, 50 – 100 mL of eluted protein solution (at ca. 1 mg mL$^{-1}$) was dialyzed against 4 L distilled water at 4 °C. The water was changed repeatedly (5 – 6X) over the course of several days. Typical yields after lyophilization ranged from 100 to 200 mg L$^{-1}$.

***Removal of endotoxin and protein sterilization.*** All batches of protein used for *in vivo* immune challenge studies (regardless of recombinant origin, i.e. *E. coli* or CHO expression) were subjected to an additional on-column endotoxin (i.e., lipopolysaccharide, LPS) removal protocol, consisting of extensive washing of the Ni-NTA agarose beads with a series of buffer systems that have been independently reported to remove the majority of endotoxin from recombinant proteins.[2-7] All buffers were prepared with endotoxin-free, double-distilled water. The column was first subjected to 5X sequential washes of 10 column volumes (CV) each of cold denaturing Buffer A (8 M urea, 100 mM $NaH_2PO_4$, 10 mM Tris, 10 mM imidazole, pH 8) supplemented with 0.1% (*v*/*v*) of the non-ionic detergent Triton X-114 (50 CV total washes). Washing was performed with the Buffer A kept cold (~4 °C) to prevent phase-partitioning of the detergent. After the last wash with detergent, the column was equilibrated with 10 CV of Buffer A alone, followed by 10 CV of a urea-free, Triton-free Buffer B (10 mM Tris, 10 mM imidazole, pH 8). This was followed by a 10 CV wash with Buffer C (10 mM Tris, 10 mM imidazole, pH 8, 60% *v*/*v* isopropanol). Alternating 10 CV washes with Buffer B and the alcoholic Buffer C were repeated 2 – 3X, followed by a 10 CV wash with Buffer B to remove the isopropanol. Finally, the resin was exchanged back into a phosphate-free denaturing Buffer D (8 M urea, 100 mM Tris, pH 8) and the protein eluted with 3 – 5 CV of Buffer D plus 250 mM imidazole (pH 8). The eluted protein was passed through a 0.2 µm disposable filter device (Stericup®, Millipore Sigma). 50 – 100 mL of eluted, filtered protein solution (at ca. 1 mg $mL^{-1}$) was dialyzed against 4 L of deionized, endotoxin-free water at 4 °C. The water was changed repeatedly (5 – 6X) over the course of several days. After the final dialysis exchange, the protein was sterilized again by filtration through a second 0.2 µm Stericup® filter, immediately flash-frozen in liquid $N_2$ and lyophilized.

***Preparation of stable CHO lines.*** Full-length XTEN genes coding for PXP and QXQ were synthesized by DNA2.0, codon-optimized for maximal expression in Chinese Hamster Ovary (CHO) cell lines. The 3' end of each CHO-optimized gene was subsequently fused to an IRES-GFP sequence ("DasherGFP", DNA2.0) via *XhoI* + *NheI* digestion. XTEN-IRES-GFP cassettes were PCR-amplified using primers containing *SapI* restriction sites, and ligated into the expression vector pD2537 (DNA2.0) with subsequent *SapI* site destruction. Each plasmid contained the secretion tag leader sequence MKWVTFISLLFLFSSAYS derived from human serum albumin at the N-terminus of each XTEN cassette (cleaved during secretion). The plasmids were amplified in *E. coli*, purified by Maxiprep with endotoxin-free wash buffer (Macherey-Nagel), complexed with ExpiFectamine CHO Reagent (ThermoFisher), and used to transfect ExpiCHO-S cells (ThermoFisher) following the manufacturer's instructions. Within two days, transfected suspension cells were transitioned into 364-well plates containing fresh ExpiCHO Expression Media supplemented with Hygromycin B (100 µg/mL), 10% FBS, and 2.5 µg/mL Fungizone

(Amphotericin B, ThermoFisher). Successfully transfected (surviving, Hygromycin resistant) clones appeared bright green by fluorescence microscopy, and were slowly expanded into T25 flasks over a period of 2 – 3 months (**Fig. S6**). Supernatants of fluorescent, expanded clones were screened by Western Blotting against 6xHis to evaluate protein expression and putative secretion into the media. The highest-expressing clones were frozen and stored in preparation for large-scale expression.

***Expression of "endotoxin free" XTEN proteins from CHO cells.*** For protein expression from stable CHO cell lines, the desired clone was expanded to 1 L in GMP-manufactured serum-free, protein-free, animal origin-free ExpiCHO Expression Medium (ThermoFisher), and incubated for 1 – 2 weeks at 37 °C shaking incubator containing 5% $CO_2$ atmosphere. Live cell density was monitored closely throughout the expression period. Cells typically exhibited a precipitous decline in viability towards the end of this period, going from high viability (>90%) to death within 12 – 24 h. Supernatant containing the desired XTEN protein was harvested during or immediately after this rapid death phase. The CHO supernatant was clarified by centrifugation (30 min at 3000 rpm followed by clarification of supernatant at 30 min at 6000 rpm at 4 °C, in a Fiberlite™ F6-6 x 1000y rotor using a Sorvall Evolution RC centrifuge), followed by passage through a 0.2 µm filter. The clarified supernatant was supplemented with 10 – 20 mM imidazole and 300 mM NaCl, and incubated with fresh Ni-NTA resin (typically 10 mL per L of culture supernatant) for at least an hour with gentle stirring at 4 °C. The resin-bound protein was then subject to the purification, endotoxin removal and sterilization procedure described above. Yields of purified protein were typically 50 – 100 mg $L^{-1}$ of clarified ExpiCHO supernatant.

***Cloning, expression and purification of αDEC-205-QX fusion antibody.*** Plasmids coding for the heavy chain (HC) and light chain (LC) of the IgG anti-mouse CD205 (DEC-205) monoclonal antibody clone NLDC-145 and isotype HC and LC control plasmids (all carrying the ampicillin resistance gene *ampR*) were the kind gift of the Nussenzweig Lab at Rockefeller University.[8] The LC plasmids were used as received. The *E. coli* codon optimized "QX" gene was installed at the 3' end of each HC gene using unique *NheI* + *NotI* restriction sites, with a $(GS)_6$ linker and 6xHis tag installed at both the N- and C-terminus of the QX coding sequence. This was done to match the corresponding sequence of the full-length QXQ protein, which also contained telechelic $(GS)_6$ and 6xHis tags. Antibody expression was performed by the Caltech Protein Expression Center

(PEC) in transiently-transfected cultures of HEK-293-6E cells. The plasmids were amplified in *E. coli* DH10B and purified by Maxiprep with endotoxin-free wash buffer (Macherey-Nagel). Transfection was performed by mixing 500 µg each of the HC and LC plasmids, complexing this mixture with polyethylenimine (PEI), and adding this mixture to the cells. Transfected 1 L cultures were grown for 1 week at 37 °C in a shaking incubator containing 5% $CO_2$ atmosphere. After 1 week, the supernatant was harvested by centrifugation and filtration. Secreted, folded antibody was purified directly from the supernatant over a 1 mL or 5 mL HisTrap™ HP IMAC column (GE Healthcare), eluting with 250 mM imidazole (this route was chosen over Protein A purification to prevent likely hydrophobic aggregation mediated by the QX domain under acidic elution conditions). Eluted antibody was dialyzed overnight against endotoxin-free, 100 mM phosphate buffer (pH 7.4), concentrated to ~0.5 mL using 10,000 NMWL Amicon Ultra Centrifugal Filter Units (Millipore Sigma), and sized over a Superose 6 Increase 10/300 GL column (GE Healthcare). To remove trace endotoxin from the column, the column was first pre-treated with 2 CV of endotoxin-free HyClone water (GE Lifesciences), 1 CV of 0.5 M NaOH in HyClone water, 2 CV of water, and finally equilibrated in 1 CV of GMP-grade HyClone DPBS (no Ca or Mg salts). Eluted antibody fractions were concentrated in DPBS to the desired concentration by centrifugal spin filtration (typically to 0.1 mg $mL^{-1}$ final concentration), filtered through a 0.2 µm filter for sterilization, and stored at 4 °C until use.

***Quantification of endotoxin levels.*** Endotoxin contamination in recombinant protein samples was assayed using the HEK-Blue™ mTLR4 reporter cell line (InvivoGen). Proteins were first dissolved in endotoxin-free 100 mM phosphate buffer, pH 7.4 at a concentration of 10 mg $mL^{-1}$ (i.e., 1% *w*/*v*). Higher concentrations induced sample gelation that could interfere with assay performance. The samples underwent serial 10-fold dilutions down to concentrations as a low as $10^{-4}$ mg $mL^{-1}$. Subsequently, 20 µL of each dilution was added to a TC-treated 96-well microplate, and a suspension of ~25,000 HEK-Blue™ cells were added to each well in 180 µL of Growth Medium (DMEM, 4.5 g $L^{-1}$ glucose, 10% *v*/*v* FBS, 50 U $mL^{-1}$ penicillin, 50 µg $mL^{-1}$ streptomycin, 100 µg $mL^{-1}$ Normocin™, 2 mM L-glutamine). The cells were incubated with protein at 37 °C in 5% $CO_2$ atmosphere for 16 h. To detect secreted alkaline phosphatase (SEAP), 20 µL of this supernatant was added to 200 µL of pre-warmed, 0.2 µm-filtered QUANTI-Blue™ (InvivoGen) and the mixture incubated for 6 – 8 h. Levels of SEAP (indicating presence of endotoxin) were quantified using a plate-reader by measuring well absorbance at 620 nm. Positive control wells were treated with serial dilutions of an LPS standard from *E. coli* strain O127:B8 (Sigma, L3129). Endotoxin mass concentrations were converted to the EU $mL^{-1}$ standard by assuming 1 EU = 0.1 ng. Negative control wells were treated with endotoxin-free water or DPBS. The limit of detection for this assay was ~0.1 EU $mL^{-1}$.

***Oscillatory shear rheology of hydrogels.*** Flow properties of elastin and XTEN hydrogels carrying native and mutant P domains were characterized by oscillatory shear rheology. Hydrogels were prepared by adding 100 µM phosphate buffer (pH 7.4) directly to lyophilized protein, and allowing the solid suspension to swell for 12 – 24 h. In all cases, the final protein mass concentration in the network was 10% *w/v*, i.e. 100 mg $mL^{-1}$. Samples were periodically mixed, allowed to set, and then centrifuged at 18,000*g* for 1 min during the swelling period to obtain homogenous networks. Oscillatory shear rheometry was performed on the swollen gels using an ARES-RFS strain-controlled rheometer (TA Instruments) equipped with a cone-and-plate geometry (25 mm diameter, gap width 50 µm). The outer edge of the plate was coated with mineral oil to minimize evaporation, and sample temperature was maintained at 25 °C or 37 °C using a circulating water batch. Strain sweeps at 10 rad $s^{-1}$ identified a linear regime between 0.1 and 10% strain. Frequency sweeps were performed at a fixed strain amplitude of 1% between 0.01 and 100 rad $s^{-1}$. The terminal elastic modulus $G_\infty$ and the characteristic network relaxation rate $\omega_c = 1 / \tau$ were determined by fitting the frequency sweep rheometry data to a Maxwell model using the following equations (non-linear curve fitting was performed in GraphPad Prism):

$$G'(\omega) = G_\infty \frac{(\omega\tau)^2}{1+(\omega\tau)^2} \qquad \text{(eq. \textbf{S1})}$$

$$G''(\omega) = G_\infty \frac{\omega\tau}{1+(\omega\tau)^2} \qquad \text{(eq. \textbf{S2})}$$

The fit parameters $G_\infty$ and $\tau$ could both be obtained independently from eq. **S1** and eq. **S2**, and were averaged to provide the data presented in **Fig. 2, Fig. S1,** and **Fig. S3**.

***Immune challenge and tolerance induction protocol.*** All animal experiments were conducted under an approved IACUC protocol in accordance with Caltech OLAR guidelines. Female BALB/cJ mice (7 weeks old) were purchased from The Jackson Laboratory (Stock No. 000651).

All mice were acclimatized to the new facilities for at least 1 week prior to the start of immune challenge or tolerance induction studies. For all *in vivo* studies involving XTEN proteins, sterile hydrogels (10% *w*/*v*) were prepared by swelling sterile-filtered, lyophilized XTEN protein in sterile endotoxin-free 100 mM phosphate buffer (pH 7.4), and loading the hydrogels into a sterile polycarbonate syringe using a heat-sterilized spatula. Mice 8 – 10 weeks old were anesthetized with 2 – 5% isoflurane for the hydrogel injection procedure. Prior to the first hydrogel injection (day 0), the mouse dorsum was carefully shaved (Wahl Pocket Pro), the hair removed and the injection site cleaned by wiping with an alcohol swap (BD Bioscience). Hydrogels were injected subcutaneously (SC) through a 22-gauge needle (50 µL per injection, corresponding to 5 mg of XTEN protein). Hydrogels were injected twice on day 0 and day 14. Positive control mice were injected twice intraperitoneally (IP) on day 0 and day 14 with 100 µL of a suspension of 1 µg $µL^{-1}$ whole Ovalbumin (OVA Grade VII, Sigma A7641) admixed to Imject® Alum (Thermo Scientific) at a 1:1 ratio (i.e., 50 µg OVA per injection). For tolerance studies, tolerized groups received 10 µg of αDEC-205-QX antibody or an isotype control antibody administered IP on day -14. For all groups, blood was collected from the mouse facial vein on day 28 into a Microtainer® Tube (BD 365967) containing a silica-based clot activator and inert polymer gel for facile clot capture. Freshly isolated blood was allowed to clot for 30 min at room temperature, then the serum was isolated from the clot by centrifuging at 1,100*g* for 10 min. Serum was diluted 1:10 into Blocking Buffer (PBS, 0.05% Tween 20, 2% BSA), and stored at –80 °C prior to analysis.

***Analysis of antibody titers by enzyme-linked immunosorbent assay (ELISA).*** Antigen-specific antibody titers were assayed in 96-well plate format by indirect ELISA. The relevant antigen (e.g. PXP, QXQ or OVA) was dissolved in Coating Buffer (100 mM phosphate buffer, pH 8) at 1 – 10 µg $mL^{-1}$, and 100 µL per well of this solution was placed onto a high-binding ELISA plate (Nunc "MaxiSorp", ThermoFisher 439454) overnight at 4 °C. The plate was then washed 5X with Wash Buffer (PBS, 0.05% Tween 20), 200 µL of Blocking Buffer (PBS, 0.05% Tween 20, 2% BSA) was added, and the plate was again incubated overnight 4 °C. Sera was removed from the freezer, thawed and diluted (typically from $1:10^2$ to $1:10^5$) in Blocking Buffer, then added to the plate at 100 µL per well. Plates were covered and incubated with the sera for 2 h at room temperature before the sera were removed, and the plate washed 5X with 200 µL of Wash Buffer. Adsorbed, antigen-specific antibodies were detected using biotin-labeled, affinity purified goat anti-mouse IgG γ-chain (SeraCare, KPL 16-18-02) diluted $1:10^3$ (0.5 µg $mL^{-1}$ final concentration) in 0.2X Blocking Buffer and incubated for 1 h at room temperature and 100 µL per well, followed by 5X washing with Wash Buffer. Peroxidase-labeled streptavidin (SeraCare KPL 14-30-00) was diluted 1:$10^2$ (5.0 µg $mL^{-1}$ final concentration) and incubated with each well for 30 min, followed by extensive washing (10X) with 200 µL Wash Buffer. Finally, the plates were developed for 2 min with 100 µL of 1X TMB (3,3',5,5'-Tetramethylbenzidine, eBioscience) substrate, followed by quenching of the color development with 100 µL of 1 M $H_2SO_4$. Absorbance was measured at 450 nm on a plate reader. Additional isotype specific, polyclonal goat anti-mouse HRP-conjugated antibodies for serum isotyping experiments were purchased from Abcam, and were used at a dilution of 1:$10^3$. To minimize precision errors introduced by rounding to the nearest (or next highest) $\log_{10}$ dilution value, antibody titers are reported as -$\log_{10}$ of the interpolated serum dilution value (in $\log_{10}$ space) yielding an $Abs_{450}$ signal of unity on the plate reader. Typically, an absorbance value of unity was intersected by the assay over the range 1:$10^2$ – 1:$10^5$, consistently providing antibody titers between 2 and 5. Extrapolation starting from the well closest to unity was required in a few instances, if a unity absorbance value was not intersected over the dilution range assayed. Wells yielding an $Abs_{450}$ value of $\leq 0.1$ at a dilution of 1:$10^2$ were assigned a titer of 1, which was taken as the limit of detection of the assay.

***Circular dichroism spectroscopy.*** Circular dichroism spectra were recorded on an Aviv Model 430 CD Spectrometer. Protein samples were prepared at 30 µM in 100 mM phosphate buffer, 0.2 µm-filtered, and scanned at room temperature between 185 nm and 300 nm in 1 nm wavelength step increments with a spectral averaging time of 5 seconds per step. The signal in millidegrees was converted to mean residue ellipticity using the formula:

$$[\theta]\ (\text{deg cm}^2\ \text{dmol}^{-1}) = \frac{\text{millidegrees}}{\text{pathlength (mm)} \times \text{concentration (M)} \times \text{number of residues}}$$

(eq. **S3**)

**Table S1. HPEPDOCK results for top 10 P and Q antigens bound in H2-I-A$^d$ .** Model ranks and relative docking energy scores for all binding poses are displayed below. Poses with the peptide incorrectly bound outside of the MHC cleft (“unbound”) are highlighted in red. The poses corresponding to the structures in **Fig. 1D** are shaded in red, blue, and yellow.

| | | *model rank and relative docking energy score* | | | | | | | | | | *average dock energy scores* | | |
|---|---|---|---|---|---|---|---|---|---|---|---|---|---|---|
| **Peptide** | **Sequence** | **1** | **2** | **3** | **4** | **5** | **6** | **7** | **8** | **9** | **10** | **avg_bound** | **avg_unbound** | **# bound** |
| $Q_{50-64}$ | LLAQQMAEIAALKMQ | -192.458 | -188.25 | -186.93 | -186.75 | -186.53 | -186.19 | -184.84 | -184.36 | -183.84 | -183.37 | -186.7 | -185.1 | 8 |
| $Q_{57-71}$ | EIAALKMQVMASDAA | -247.39 | -233.82 | -209.78 | -199.25 | -198.53 | -195.26 | -193.76 | -190.36 | -190.11 | -188.41 | -204.7 | - | 10 |
| $Q_{49-63}$ | ALLAQQMAEIAALKM | -211.231 | -202.42 | -200.52 | -198.71 | -193.27 | -192.09 | -190.46 | -190.08 | -189.42 | -187.91 | -195.6 | - | 10 |
| $Q_{51-65}$ | LAQQMAEIAALKMQV | -220.386 | -207.56 | -204.22 | -204.15 | -199.08 | -197.73 | -194.89 | -193.31 | -192.39 | -190.72 | -200.4 | - | 10 |
| $Q_{53-67}$ | QQMAEIAALKMQVMA | -218.427 | -209.12 | -206.72 | -200.61 | -199.10 | -191.85 | -191.47 | -187.78 | -185.33 | -185.13 | -197.6 | - | 10 |
| $Q_{52-66}$ | AQQMAEIAALKMQVM | -207.648 | -203.28 | -202.03 | -189.28 | -189.25 | -188.36 | -186.26 | -185.11 | -184.90 | -184.81 | -191.8 | -193.4 | 9 |
| $Q_{48-62}$ | RALLAQQMAEIAALK | -236.424 | -222.77 | -217.34 | -215.35 | -203.99 | -201.41 | -201.06 | -199.87 | -199.41 | -195.22 | -211.5 | -200.4 | 8 |
| $Q_{47-61}$ | VRALLAQQMAEIAAL | -233.10 | -228.81 | -222.86 | -218.71 | -218.31 | -208.09 | -207.62 | -199.62 | -194.86 | -194.36 | -212.6 | - | 10 |
| $Q_{38-52}$ | MATNMALADVRALLA | -223.173 | -216.51 | -213.87 | -208.54 | -202.91 | -200.38 | -200.29 | -199.83 | -199.44 | -196.99 | -205.0 | -216.5 | 9 |
| $Q_{39-53}$ | ATNMALADVRALLAQ | -217.116 | -209.58 | -206.22 | -197.50 | -196.47 | -193.87 | -193.49 | -192.46 | -192.08 | -191.45 | -199.8 | -192.5 | 9 |
| | **avg_rank_1** | -220.735 | | | | | | | | | | **-200.6** | **-197.6** | |

| | | | | | | | | | | | | | | |
|---|---|---|---|---|---|---|---|---|---|---|---|---|---|---|
| $P_{35-49}$ | RELQETNAALQDVRE | -175.89 | -175.56 | -173.68 | -173.46 | -170.67 | -166.45 | -165.49 | -165.16 | -164.93 | -164.37 | -169.4 | -170.1 | 8 |
| $P_{34-48}$ | LRELQETNAALQDVR | -193.15 | -192.19 | -190.69 | -190.61 | -184.11 | -181.75 | -181.33 | -178.36 | -175.81 | -173.90 | -184.8 | -178.4 | 9 |
| $P_{49-63}$ | ELLRQQVKEITFLKN | -207.57 | -182.41 | -180.05 | -179.22 | -177.10 | -175.97 | -173.95 | -172.62 | -167.54 | -167.30 | -181.8 | -170.3 | 7 |
| $P_{47-61}$ | VRELLRQQVKEITFL | -216.17 | -196.07 | -195.34 | -193.22 | -188.26 | -187.87 | -187.54 | -187.09 | -187.04 | -185.30 | -191.1 | -194.3 | 6 |
| $P_{36-50}$ | ELQETNAALQDVREL | -200.28 | -196.26 | -190.06 | -180.53 | -180.38 | -178.71 | -173.82 | -171.93 | -168.19 | -166.59 | -181.6 | -171.9 | 9 |
| $P_{33-47}$ | MLRELQETNAALQDV | -213.45 | -185.18 | -183.53 | -176.42 | -175.34 | -175.25 | -174.91 | -174.91 | -174.59 | -173.49 | -180.7 | - | 10 |
| $P_{41-55}$ | NAALQDVRELLRQQV | -183.50 | -180.93 | -178.24 | -177.55 | -173.53 | -171.73 | -171.47 | -170.61 | -169.13 | -168.45 | -174.9 | -173.5 | 7 |
| $P_{48-62}$ | RELLRQQVKEITFLK | -199.36 | -198.85 | -190.83 | -183.70 | -183.29 | -182.42 | -177.75 | -177.62 | -177.31 | -177.06 | -184.9 | -183.7 | 9 |
| $P_{42-56}$ | AALQDVRELLRQQVK | -204.69 | -196.79 | -187.66 | -187.07 | -182.73 | -180.31 | -177.79 | -171.30 | -170.77 | -167.21 | -185.6 | -178.2 | 6 |
| $P_{50-64}$ | LLRQQVKEITFLKNT | -219.93 | -207.12 | -206.42 | -205.41 | -204.59 | -201.48 | -198.80 | -196.17 | -194.93 | -188.68 | -202.4 | - | 10 |
| | **avg_rank_1** | -201.40 | | | | | | | | | | **-183.7** | **-177.6** | |

| | | | | | | | | | | | | | | |
|---|---|---|---|---|---|---|---|---|---|---|---|---|---|---|
| $OVA_{323-339}$ | **rank_1** | -232.63 | -230.98 | -219.58 | -214.92 | -214.89 | -213.08 | -211.85 | -211.66 | -211.21 | -211.20 | **-217.2** | - | 10 |

**Table S2. Comparative immunogenicity analysis of other coiled-coil domains.** Sequences of other coiled-coil forming domains taken from the literature[9-18] were analyzed for their ability to interact with murine MHC Class II (H2-I-$A^d$). Reported below are the average Consensus Percentile Rank and NN Align $IC_{50}$ values for the top five peptide:MHC binding registers from each sequence, as predicted by the IEDB. The binding predictions for P, Q, and OVA reported in **Fig. 1** and **Fig. S3** differ slightly from values reported below, as the latter predictions were generated with a newer version of the IEDB web server (v2023).

| **Coil Name / Sequence** | **$IC_{50}$ (μM)** | **log [%ile rank]** | **Reference (DOI)** |
|---|---|---|---|
| COMPcc (P)<br>APQMLRELQETNAALQDVRELLRQQVKEITFLKNTVMESDAS | 3.328 | 1.51 | 10.1021/bi900534r |
| de novo tetramer (A)<br>SGDLENEVAQLEREVRSLEDEAAELEQKVSRLKNEIEDLKAE | 13.149 | 1.64 | 10.1038/nmat1573 |
| de novo phenylalanine pentamer ($F_V$)<br>SSNAKFDQFSSDFQTFNAKFDQFSNDFNAFRSDFQAFKDDFARFNQRFDNFATKYR | 16.868 | 1.86 | 10.1016/j.jmb.2006.05.063 |
| de novo phenylalanine tetramer ($F_{IV}$)<br>SSNAKFDQFSSDFQTFNAKFDQFSNDMNAFRSDFQAFKDDFARFNQRFDNFATKYR | 9.796 | 1.84 | 10.1016/j.jmb.2006.05.063 |
| tetrabrachion tetramer (T)<br>GSIINETADDIVYRLTVIIDDRYESLKNLITLRADRLEMIINDNVSTILASG | 0.176 | 0.76 | 10.1038/79006 |
| influenza A homodimer ($I_\alpha$)<br>QNRNGKWREQLGQKFEEIRWLIEEVRHRLKITENSFEQITFMQALQLLLEVEQEIRTF | 0.551 | 0.88 | 10.1093/emboj/cdg449 |
| influenza B homodimer ($I_\beta$)<br>QKRETIRLVTEELYLLSKRIDDNILFHKTVIANSSIIADMVVSLSLLETLYEMKDVIEVY | 0.171 | 0.71 | 10.1093/emboj/cdg449 |
| sendai virus phosphoprotein tetramer (S)<br>YAEMTFNVCGLILSAEKSSARKVDENKQLLKQIQESVESFRDIYKRFSEYQKEQNSLLMSNLSTL | 1.485 | 1.15 | 10.1038/79013 |
| tetramer from mSNAP-23 (N)<br>STRRILGLAIESQDAGIKTITMLDEQGEQLNRIEEGMDQINKDMREAEKTLTEL | 1.923 | 1.33 | 10.1074/jbc.m210483200 |
| tetrameric LacI repressor (L)<br>PRALADSLMQLARQVSRLESGQ | 1.321 | 1.02 | 10.1002/pro.5560040803 |
| GCN4-derived tetramer ($G_{IV}$)<br>MKVKQLVDKVEELLSKNYHLVNEVARLVKLVGER | 2.647 | 1.36 | 10.1021/bi061914m |
| GCN4-derived heptamer ($G_{VII}$)<br>MKVKQLADAVEELASANYHLANAVARLAKAVGER | 0.444 | 0.71 | 10.1073/pnas.0604871103 |
| **evolved from P** (Q)<br>AEEMLRELMATNMALADVRALLAQQMAEIAALKMQVMASDAA | 0.030 | -0.72 | ***this work*** |
| ovalbumin (OVA)<br>MGSIGAASMEFCFDVFKELKVHHANENIFYCPIAIMSALAMVYL... | 0.047 | -0.29 | uniprot A0A411G5W6 |

**Table S3. Amino acid sequences of artificial proteins, hydrogels, and antibody fusions.** The coiled-coil forming domains P and Q were used to prepare hydrogel forming proteins flanking a central flexible X domain (derived from nonimmunogenic XTEN), as well as antibody fusions carrying a C-terminal "QX" tail encoded on the heavy chain.

| Domain Name | Length | MW (kDa) | Amino Acid Sequence |
|---|---|---|---|
| P | 42 | - | APQMLRELQETNAALQDVRELLRQQVKEITFLKNTVMESDAS |
| Q | 42 | - | AEEMLRELMATNMALADVRALLAQQMAEIAALKMQVMASDAA |
| X | 144 | - | GSPAGSPTSTEEGTSESATPESGPGTSTEPSEGSAPGSPAGSPTSTEEGTSTE<br>PSEGSAPGTSTEPSEGSAPGTSESATPESGPGSEPATSGSETPGSEPATSGSE<br>TPGSPAGSPTSTEEGTSESATPESGPGTSTEPSEGSAP |
| **Hydrogel Proteins** | | | |
| PXP | 307 | 29.9 | MRGSH$_6$GSVD(GS)$_6$G{**P**}(GS)$_6$LD{**X**}LD(GS)$_6$G{**P**}(GS)$_6$LEH$_6$KLN* |
| QXQ | 307 | 29.3 | MRGSH$_6$GSVD(GS)$_6$G{**Q**}(GS)$_6$LD{**X**}LD(GS)$_6$G{**Q**}(GS)$_6$LEH$_6$KLN* |
| QX | 238 | 22.8 | MRGSH$_6$GSVD(GS)$_6$G{**Q**}(GS)$_6$LD{**X**}LEH$_6$KLN* |
| X | 169 | 16.3 | MRGSH$_6$GSVD{**X**}LEH$_6$KLN* |
| **Antibody Fusions** | | | |
| αDEC-QX | 738 | 77.4 | MGWSCIILFLVATATGVHSEVKLLESGGGLVQPGGSLRLSCAASGFTFNDFYM<br>NWIRQPPGQAPEWLGVIRNKGNGYTTEVNTSVKGRFTISRDNTQNILYLQMNS<br>LRAEDTAIYYCARGGPYYYSGDDAPYWGQGVMVTVSSATTKGPSVYPLAPGSA<br>AQTNSMVTLGCLVKGYFPEPVTVTWNSGSLSSGVHTFPAVLQSDLYTLSSSVT<br>VPSSTWPSETVTCNVAHPASSTKVDKKIVPRDCGCKPCICTVPEVSSVFIFPP<br>KPKDVLTITLTPKVTCVVVAISKDDPEVQFSWFVDDVEVHTAQTQPREEQFNS<br>TFRSVSELPIMHQDWLNGKEFKCRVNSAAFPAPIEKTISKTKGRPKAPQVYTI<br>PPPKEQMAKDKVSLTCMITDFFPEDITVEWQWNGQPAENYKNTQPIMDTDGSY<br>FVYSKLNVQKSNWEAGNTFTCSVLHEGLHNHHTEKSLSHSPGKASDMAKKETV<br>WRLEEFGRFMRGSH$_6$GSVD(GS)$_6$G{**Q**}(GS)$_6${**X**}LEH$_6$KLN* |
| αDEC-κ | 233 | 25.7 | MGWSCIILFLVATATGVHSDIQMTQSPSFLSTSLGNSITITCHASQNIKGWLA<br>WYQQKSGNAPQLLIYKASSLQSGVPSRFSGSGSGTDYIFTISNLQPEDIATYY<br>CQHYQSFPWTFGGGTKLELKRADAAPTVSIFPPSSEQLTSGGASVVCFLNNFY<br>PKDINVKWKIDGSERQNGVLNSWTDQDSKDSTYSMSSTLTLTKDEYERHNSYT<br>CEATHKTSTSPIVKSFNRNEC* |
| ISO-QX | 733 | 77.0 | MGWSCIILFLVATATGVHSQVQLKESGPGLVQPSQTLSLTCTVSGFSLISYHV<br>TWVRQPPGKSLVWMGTIWTGGGRNYNSAEQSRLSISRDTSKSQVFLKMNSLQP<br>EDTGTYYCARHRGGYNYGFDYWGQGVMVTVSSATTKGPSVYPLAPGSAAQTNS<br>MVTLGCLVKGYFPEPVTVTWNSGSLSSGVHTFPAVLQSDLYTLSSSVTVPSST<br>WPSETVTCNVAHPASSTKVDKKIVPRDCGCKPCICTVPEVSSVFIFPPKPKDV<br>LTITLTPKVTCVVVAISKDDPEVQFSWFVDDVEVHTAQTQPREEQFNSTFRSV<br>SELPIMHQDWLNGKEFKCRVNSAAFPAPIEKTISKTKGRPKAPQVYTIPPPKE<br>QMAKDKVSLTCMITDFFPEDITVEWQWNGQPAENYKNTQPIMDTDGSYFVYSK<br>LNVQKSNWEAGNTFTCSVLHEGLHNHHTEKSLSHSPGKASDMAKKETVWRLEE<br>FGRFMRGSH$_6$GSVD(GS)$_6$G{**Q**}(GS)$_6${**X**}LEH$_6$KLN* |
| ISO-κ | 248 | 27.1 | MGWSCIILFLVATATGVHSDIQMTQSPSLLSASVGDRVTLNCKASQNINKNLD<br>WYQQKLGEAPKVLIYYTDNLQTGFSSRFSGSGSGTDYTLTISSLQPEDVATYY<br>CYQYNSGPTFGPGTKLELKRADAAPTVSIFPPSSEQLTSGGASVVCFLNNFYP<br>KDINVKWKIDGSERQNGVLNSWTDQDSKDSTYSMSSTLTLTKDEYERHNSYTC<br>EATHKTSTSPIVKSFNRNECASGGGGGGDYKDDDDK* |

**Table S4. Plasmid sequences of artificial proteins, hydrogels, and antibody fusions.** The nucleotide sequence encoding the secretion tag leader peptide MKWVTFISLLFLFSSAYS (derived from human serum albumin) present at the 5' end of the CHO clones is underlined.

| Hydrogel Proteins | *Host* (Plasmid) | Coding Sequence |
|---|---|---|
| PXP | *E. coli* (pQE-80L) | atgagaggatcgcatcaccatcaccatcacgGATCCGTCGACGGCTCAGGTTCGGGTAGTGGGAGCGGCTCTGGCAGCGGTGCGCCGCAAATGCT<br>GCGTGAACTGCAGGAAACCAATGCCGCGCTTCAGGATGTGCGGGAATTGCTTCGTCAACAGGTCAAGGAGATAACGTTCTTGAAGAACACCGTCA<br>TGGAGTCGGATGCGTCCGGATCTGGCTCCGGAAGCGGAAGTGGTTCTGGTAGCCTCGACGGCTCCCCGGCTGGCAGCCCGACCAGCACTGAAGAG<br>GGCACGAGCGAGTCGGCGACCCCGGAGTCTGGTCCGGGCACCTCCACCGAACCGTCTGAGGGCAGCGCACCGGGTAGCCCGGCCGGTAGCCCTAC<br>CAGCACCGAAGAGGGTACCAGCACGGAACCGAGCGAAGGCTCGGCACCGGGTACGAGCACCGAGCCGTCCGAGGGTTCCGCGCCAGGTACCAGCG<br>AGAGCGCAACGCCGGAGTCCGGTCCGGGCAGCGAACCAGCGACCAGCGGCAGCGAAACGCCGGGTTCAGAGCCGGCGACGAGCGGTAGCGAGACT<br>CCGGGCAGCCCGGCTGGTAGCCCGACGTCCACCGAAGAAGGCACCAGCGAAAGCGCCACCCCGGAGAGCGGTCCTGGTACGTCTACCGAGCCATC<br>TGAGGGTAGCGCGCCGCTCGACGGCTCAGGTTCGGGTAGTGGGAGCGGCTCTGGCAGCGGTGCGCCGCAAATGCTGCGTGAACTGCAGGAAACCA<br>ATGCCGCGCTTCAGGATGTGCGGGAATTGCTTCGTCAACAGGTCAAGGAGATAACGTTCTTGAAGAACACCGTCATGGAGTCGGATGCGTCCGGA<br>TCTGGCTCCGGAAGCGGAAGTGGTTCTGGTAGCCTCGAGCATCACCATCACCATCACAagcttaattag |
| QXQ | *E. coli* (pQE-80L) | atgagaggatcgcatcaccatcaccatcacgGATCCGTCGACGGCTCAGGTTCGGGTAGTGGGAGCGGCTCTGGCAGCGGTGCGGAGGAAATGCT<br>GCGTGAACTGATGGCAACCAATATGGCGCTTGCGGATGTGCGGGCATTGCTTGCTCAACAGATGGCGGAGATAGCGGCCTTGAAGATGCAAGTCA<br>TGGCGTCGGATGCGGCCGGATCTGGCTCCGGAAGCGGAAGTGGTTCTGGTAGCCTCGACGGCTCCCCGGCTGGCAGCCCGACCAGCACTGAAGAG<br>GGCACGAGCGAGTCGGCGACCCCGGAGTCTGGTCCGGGCACCTCCACCGAACCGTCTGAGGGCAGCGCACCGGGTAGCCCGGCCGGTAGCCCTAC<br>CAGCACCGAAGAGGGTACCAGCACGGAACCGAGCGAAGGCTCGGCACCGGGTACGAGCACCGAGCCGTCCGAGGGTTCCGCGCCAGGTACCAGCG<br>AGAGCGCAACGCCGGAGTCCGGTCCGGGCAGCGAACCAGCGACCAGCGGCAGCGAAACGCCGGGTTCAGAGCCGGCGACGAGCGGTAGCGAGACT<br>CCGGGCAGCCCGGCTGGTAGCCCGACGTCCACCGAAGAAGGCACCAGCGAAAGCGCCACCCCGGAGAGCGGTCCTGGTACGTCTACCGAGCCATC<br>TGAGGGTAGCGCGCCGCTCGACGGCTCAGGTTCGGGTAGTGGGAGCGGCTCTGGCAGCGGTGCGGAGGAAATGCTGCGTGAACTGATGGCAACCA<br>ATATGGCGCTTGCGGATGTGCGGGCATTGCTTGCTCAACAGATGGCGGAGATAGCGGCCTTGAAGATGCAAGTCATGGCGTCGGATGCGGCCGGA<br>TCTGGCTCCGGAAGCGGAAGTGGTTCTGGTAGCCTCGAGCATCACCATCACCATCACAagcttaattag |
| QX | *E. coli* (pQE-80L) | atgagaggatcgcatcaccatcaccatcacgGATCCGTCGACGGCTCAGGTTCGGGTAGTGGGAGCGGCTCTGGCAGCGGTGCGGAGGAAATGCT<br>GCGTGAACTGATGGCAACCAATATGGCGCTTGCGGATGTGCGGGCATTGCTTGCTCAACAGATGGCGGAGATAGCGGCCTTGAAGATGCAAGTCA<br>TGGCGTCGGATGCGGCCGGATCTGGCTCCGGAAGCGGAAGTGGTTCTGGTAGCCTCGACGGCTCCCCGGCTGGCAGCCCGACCAGCACTGAAGAG<br>GGCACGAGCGAGTCGGCGACCCCGGAGTCTGGTCCGGGCACCTCCACCGAACCGTCTGAGGGCAGCGCACCGGGTAGCCCGGCCGGTAGCCCTAC<br>CAGCACCGAAGAGGGTACCAGCACGGAACCGAGCGAAGGCTCGGCACCGGGTACGAGCACCGAGCCGTCCGAGGGTTCCGCGCCAGGTACCAGCG<br>AGAGCGCAACGCCGGAGTCCGGTCCGGGCAGCGAACCAGCGACCAGCGGCAGCGAAACGCCGGGTTCAGAGCCGGCGACGAGCGGTAGCGAGACT<br>CCGGGCAGCCCGGCTGGTAGCCCGACGTCCACCGAAGAAGGCACCAGCGAAAGCGCCACCCCGGAGAGCGGTCCTGGTACGTCTACCGAGCCATC<br>TGAGGGTAGCGCGCCGCTCGAGCATCATCACCACCACCATAagcttaattag |
| X | *E. coli* (pQE-80L) | atgagaggatcgcatcaccatcaccatcacgGATCCGTCGACGGCTCCCCGGCTGGCAGCCCGACCAGCACTGAAGAGGGCACGAGCGAGTCGGC<br>GACCCCGGAGTCTGGTCCGGGCACCTCCACCGAACCGTCTGAGGGCAGCGCACCGGGTAGCCCGGCCGGTAGCCCTACCAGCACCGAAGAGGGTA<br>CCAGCACGGAACCGAGCGAAGGCTCGGCACCGGGTACGAGCACCGAGCCGTCCGAGGGTTCCGCGCCAGGTACCAGCGAGAGCGCAACGCCGGAG<br>TCCGGTCCGGGCAGCGAACCAGCGACCAGCGGCAGCGAAACGCCGGGTTCAGAGCCGGCGACGAGCGGTAGCGAGACTCCGGGCAGCCCGGCTGG<br>TAGCCCGACGTCCACCGAAGAAGGCACCAGCGAAAGCGCCACCCCGGAGAGCGGTCCTGGTACGTCTACCGAGCCATCTGAGGGTAGCGCGCCGC<br>TCGAGCATCATCACCACCACCATAagcttaattag |
| PXP | *CHO* (pD2537) | ATGaagtgggtgaccttcatctccCTGCTGTTCCTGTTCTCCTCCGCCTACTCCATGCGCGGCTCACACCATCACCACCACCACGGATCCGTCGA<br>CGGCTCAGGCAGCGGTAGCGGCTCCGGAAGCGGTTCAGGAGCTCCACAAATGCTCCGGGAGCTTCAGGAAACCAACGCGGCTCTGCAGGATGTGC<br>GCGAGCTGCTGAGGCAGCAAGTCAAAGAGATCACCTTCCTGAAGAACACCGTGATGGAGTCGGACGCCTCCGGTTCCGGCTCAGGATCGGGAAGC<br>GGATCTGGCAGCCTGGATGGCTCCCCCGCCGGGTCCCCTACGTCAACCGAAGAAGGAACCTCCGAATCCGCCACCCCCGAATCCGGTCCAGGAAC<br>CAGCACCGAACCTTCCGAGGGCAGCGCCCCGGGATCGCCTGCGGGGAGCCCGACTTCCACTGAGGAAGGCACATCCACTGAGCCGTCCGAGGGAT<br>CAGCCCCCGGTACCAGCACTGAACCGAGCGAAGGATCGGCCCCCGGCACCTCCGAGTCTGCCACTCCGGAGTCCGGACCGGGGTCCGAGCCTGCA<br>ACTTCCGGGTCCGAAACTCCAGGATCGGAGCCTGCCACTAGCGGCAGCGAAACCCCGGGATCACCGGCCGGCTCCCCCACCTCGACCGAGGAAGG<br>GACCTCCGAGAGCGCGACCCCTGAGTCGGGGCCCGGAACCTCCACCGAACCCAGCGAAGGGTCCGCCCCGCTCGATGGGTCCGGATCAGGGTCGG<br>GATCGGGCAGCGGCTCGGGCGCTCCCCAGATGCTGAGAGAACTTCAGGAGACTAACGCGGCGCTGCAGGACGTGCGCGAACTGCTGCGGCAACAG<br>GTCAAGGAGATTACCTTCTTGAAGAACACTGTGATGGAATCGGACGCCAGCGGTTCGGGCTCGGGCTCCGGCTCCGGCTCGGGAAGCCTCGAGCA<br>CCACCATCATCACCACAAGCTTAACTGA |
| QXQ | *CHO* (pD2537) | ATGaagtgggtgaccttcatctccCTGCTGTTCCTGTTCTCCTCCGCCTACTCCATGCGCGGCTCACACCATCACCACCACCACGGATCCGTCGA<br>CGGCTCGGGATCGGGCTCCGGAAGCGGCTCCGGTTCAGGCGCCGAGGAAATGCTGAGAGAGCTCATGGCGACCAACATGGCGCTGGCAGACGTGC<br>GGGCGCTGCTGGCCCAGCAGATGGCCGAAATTGCGGCCCTCAAGATGCAAGTGATGGCCTCCGACGCGGCCGGCTCCGGGTCTGGCTCCGGATCG<br>GGATCAGGTTCCCTCGATGGGTCCCCAGCTGGTTCCCCTACGTCAACCGAAGAAGGAACCTCCGAATCCGCCACCCCCGAATCCGGTCCAGGAAC<br>CAGCACCGAACCTTCCGAGGGCAGCGCCCCGGGATCGCCTGCGGGGAGCCCGACTTCCACTGAGGAAGGCACATCCACTGAGCCGTCCGAGGGAT<br>CAGCCCCCGGTACCAGCACTGAACCGAGCGAAGGATCGGCCCCCGGCACCTCCGAGTCTGCCACTCCGGAGTCCGGACCGGGGTCCGAGCCTGCA<br>ACTTCCGGGTCCGAAACTCCAGGATCGGAGCCTGCCACTAGCGGCAGCGAAACCCCGGGATCACCGGCCGGCTCCCCCACCTCGACCGAGGAAGG<br>GACCTCCGAGAGCGCGACCCCTGAGTCGGGGCCCGGAACCTCCACCGAACCCAGCGAAGGCTCGGCTCCCCTGGACGGCTCCGGATCAGGGTCGG<br>GCAGCGGCAGCGGTTCCGGCGCTGAAGAGATGCTGAGGGAGCTGATGGCCACCAACATGGCCCTGGCCGATGTGCGGGCCTTGCTGGCTCAGCAG<br>ATGGCCGAGATCGCCGCACTGAAGATGCAAGTCATGGCATCCGACGCTGCCGGATCAGGAAGCGGCTCCGGTTCGGGGTCCGGTAGCCTCGAGCA<br>CCACCATCATCACCACAAGCTTAACTGA |

… **Table S4** continued:

| Antibody Fusions | *Host* (Plasmid) | Coding Sequence |
|---|---|---|
| αDEC-QX | *HEK293* (pROCK) | ATGGGATGGTCATGTATCATCCTTTTTCTAGTAGCAACTGCAACTGGAGTACATTCAGAGGTGAAGCTGTTGGAATCTGGAGGAGGTTTGGTACA<br>GCCGGGGGGTTCTCTGAGACTCTCCTGTGCAGCTTCTGGATTCACCTTCAATGATTTCTACATGAACTGGATCCGCCAGCCTCCAGGGCAGGCAC<br>CTGAGTGGTTGGGTGTTATTAGAAACAAAGGTAATGGTTACACAACAGAGGTCAATACATCTGTGAAGGGGCGGTTCACCATCTCCAGAGATAAT<br>ACCCAAAACATCCTCTATCTTCAAATGAACAGCCTGAGAGCTGAGGACACCGCCATTTACTACTGTGCAAGAGGCGGTCCTTATTACTACAGTGG<br>TGACGACGCCCCTTACTGGGGCCAAGGAGTCATGGTCACAGTCTCCTCAGCCACCACCAAGGGCCCATCTGTCTATCCACTGGCCCCTGGATCTG<br>CTGCCCAAACTAACTCCATGGTGACCCTGGGATGCCTGGTCAAGGGCTATTTCCCTGAGCCAGTGACAGTGACCTGGAACTCTGGATCCCTGTCC<br>AGCGGTGTGCACACCTTCCCAGCTGTCCTGCAGTCTGACCTCTACACTCTGAGCAGCTCAGTGACTGTCCCCTCCAGCACCTGGCCCAGCGAGAC<br>CGTCACCTGCAACGTTGCCCACCCGGCCAGCAGCACCAAGGTGGACAAGAAAATTGTGCCCAGGGATTGTGGTTGTAAGCCTTGCATATGTACAG<br>TCCCAGAAGTATCATCTGTCTTCATCTTCCCCCCAAAGCCCAAGGATGTGCTCACCATTACTCTGACTCCTAAGGTCACGTGTGTTGTGGTAGCA<br>ATCAGCAAGGATGATCCCGAGGTCCAGTTCAGCTGGTTTGTAGATGATGTGGAGGTGCACACAGCTCAGACGCAACCCCGGGAGGAGCAGTTCAA<br>CAGCACTTTCCGCTCAGTCAGTGAACTTCCCATCATGCACCAGGACTGGCTCAATGGCAAGGAGTTCAAATGCAGGGTCAACAGTGCAGCTTTCC<br>CTGCCCCCATCGAGAAAACCATCTCCAAAACCAAAGGCAGACCGAAGGCTCCACAGGTGTACACCATTCCACCTCCCAAGGAGCAGATGGCCAAG<br>GATAAAGTCAGTCTGACCTGCATGATAACAGACTTCTTCCCTGAAGACATTACTGTGGAGTGGCAGTGGAATGGGCAGCCAGCGGAGAACTACAA<br>GAACACTCAGCCCATCATGGACACAGATGGCTCTTACTTCGTCTACAGCAAGCTCAATGTGCAGAAGAGCAACTGGGAGGCAGGAAATACTTTCA<br>CCTGCTCTGTGTTACATGAGGGCCTGCACAACCACCATACTGAGAAGAGCCTCTCCCACTCTCCTGGTAAAgctagcgacatggccaagaaggag<br>acagtctggaggctcgaAgagttcggtaggttcATGAGAGGATCGCATCACCATCACCATCACGGATCCGTCGACGGGTCGGGATCGGGCTCCGG<br>TTCGGGCTCGGGTTCAGGAGCCGAGGAGATGCTCAGAGAACTGATGGCCACCAACATGGCCTTGGCTGATGTGCGGGCCCTGCTGGCGCAGCAGA<br>TGGCCGAAATCGCGGCCCTGAAGATGCAAGTGATGGCATCAGATGCAGCGGGCTCCGGCTCCGGCAGCGGTAGCGGGTCCGGATCTCTGGACGGA<br>AGCCCAGCTGGTTCCCCTACGTCAACCGAAGAAGGAACCTCCGAATCCGCCACCCCCGAATCCGGTCCAGGAACCAGCACCGAACCTTCCGAGGG<br>CAGCGCCCCGGGATCGCCTGCGGGGAGCCCGACTTCCACTGAGGAAGGCACATCCACTGAGCCGTCCGAGGGATCAGCCCCCGGTACCAGCACTG<br>AACCGAGCGAAGGATCGGCCCCCGGCACCTCCGAGTCTGCCACTCCGGAGTCCGGACCGGGGTCCGAGCCTGCAACTTCCGGGTCCGAAACTCCA<br>GGATCGGAGCCTGCCACTAGCGGCAGCGAAACCCCGGGATCACCGGCCGGCTCCCCCACCTCGACCGAGGAAGGGACCTCCGAGAGCGCGACCCC<br>TGAGTCGGGGCCCGGAACCTCCACCGAACCCAGCGAAGGCTCGGCTCCCCTGGACGGCTCCGGCTCCGGGAGCGGAAGCGGATCAGGCTCCCTCG<br>AGCATCATCACCACCACCATAAGCTTAATTAG |
| αDEC-κ | HEK293 (pROCK) | ATGGGATGGTCATGTATCATCCTTTTTCTAGTAGCAACTGCAACTGGAGTACATTCAGACATCCAGATGACACAGTCTCCGTCATTTCTGTCTAC<br>ATCTCTTGGAAACAGCATCACCATCACTTGCCATGCCAGTCAGAACATCAAGGGTTGGTTAGCCTGGTACCAACAAAAGTCAGGGAATGCTCCTC<br>AACTGTTGATTTATAAGGCATCTAGCCTGCAATCAGGGGTTCCATCAAGATTCAGTGGCAGTGGATCTGGAACAGATTATATTTTCACTATCAGC<br>AACCTACAGCCTGAAGATATTGCCACTTATTACTGTCAGCATTATCAAAGCTTTCCGTGGACGTTCGGTGGAGGCACCAAGCTGGAATTGAAACG<br>GGCTGATGCTGCACCAACTGTATCCATCTTCCCACCATCCAGTGAGCAGTTAACATCTGGAGGTGCCTCAGTCGTGTGCTTCTTGAACAACTTCT<br>ACCCCAAAGACATCAATGTCAAGTGGAAGATTGATGGCAGTGAACGACAAAATGGCGTCCTGAACAGTTGGACTGATCAGGACAGCAAAGACAGC<br>ACCTACAGCATGAGCAGCACCCTCACGTTGACCAAGGACGAGTATGAACGACATAACAGCTATACCTGTGAGGCCACTCACAAGACATCAACTTC<br>ACCCATTGTCAAGAGCTTCAACAGGAATGAGTGTTGA |
| ISO-QX | *HEK293* (pROCK) | ATGGGATGGTCATGTATCATCCTTTTTCTAGTAGCAACTGCAACTGGAGTACATTCACAGGTGCAGCTGAAAGAGTCAGGACCTGGTCTGGTGCA<br>GCCCTCACAGACCCTGTCTCTCACCTGCACTGTCTCTGGGTTCTCACTAATCAGCTATCATGTAACCTGGGTTCGCCAGCCTCCTGGAAAGAGTC<br>TGGTGTGGATGGGAACAATATGGACTGGTGGAGGTAGAAATTATAATTCGGCTGAACAATCCCGACTGAGCATCAGCCGGGACACCTCCAAGAGC<br>CAAGTTTTCTTAAAAATGAACAGTCTGCAACCTGAAGACACAGGCACTTACTACTGTGCCAGACATCGAGGGGGGTATAACTACGGCTTTGATTA<br>CTGGGGCCAAGGAGTCATGGTCACAGTCTCCTCAGCCACCACCAAGGGCCCATCTGTCTATCCACTGGCCCCTGGATCTGCTGCCCAAACTAACT<br>CCATGGTGACCCTGGGATGCCTGGTCAAGGGCTATTTCCCTGAGCCAGTGACAGTGACCTGGAACTCTGGATCCCTGTCCAGCGGTGTGCACACC<br>TTCCCAGCTGTCCTGCAGTCTGACCTCTACACTCTGAGCAGCTCAGTGACTGTCCCCTCCAGCACCTGGCCCAGCGAGACCGTCACCTGCAACGT<br>TGCCCACCCGGCCAGCAGCACCAAGGTGGACAAGAAAATTGTGCCCAGGGATTGTGGTTGTAAGCCTTGCATATGTACAGTCCCAGAAGTATCAT<br>CTGTCTTCATCTTCCCCCCAAAGCCCAAGGATGTGCTCACCATTACTCTGACTCCTAAGGTCACGTGTGTTGTGGTAGCAATCAGCAAGGATGAT<br>CCCGAGGTCCAGTTCAGCTGGTTTGTAGATGATGTGGAGGTGCACACAGCTCAGACGCAACCCCGGGAGGAGCAGTTCAACAGCACTTTCCGCTC<br>AGTCAGTGAACTTCCCATCATGCACCAGGACTGGCTCAATGGCAAGGAGTTCAAATGCAGGGTCAACAGTGCAGCTTTCCCTGCCCCCATCGAGA<br>AAACCATCTCCAAAACCAAAGGCAGACCGAAGGCTCCACAGGTGTACACCATTCCACCTCCCAAGGAGCAGATGGCCAAGGATAAAGTCAGTCTG<br>ACCTGCATGATAACAGACTTCTTCCCTGAAGACATTACTGTGGAGTGGCAGTGGAATGGGCAGCCAGCGGAGAACTACAAGAACACTCAGCCCAT<br>CATGGACACAGATGGCTCTTACTTCGTCTACAGCAAGCTCAATGTGCAGAAGAGCAACTGGGAGGCAGGAAATACTTTCACCTGCTCTGTGTTAC<br>ATGAGGGCCTGCACAACCACCATACTGAGAAGAGCCTCTCCCACTCTCCTGGTAAAgctagcgacatggccaagaaggagacagtctggaggctc<br>gaAgagttcggtaggttcATGAGAGGATCGCATCACCATCACCATCACGGATCCGTCGACGGGTCGGGATCGGGCTCCGGTTCGGGCTCGGGTTC<br>AGGAGCCGAGGAGATGCTCAGAGAACTGATGGCCACCAACATGGCCTTGGCTGATGTGCGGGCCCTGCTGGCGCAGCAGATGGCCGAAATCGCGG<br>CCCTGAAGATGCAAGTGATGGCATCAGATGCAGCGGGCTCCGGCTCCGGCAGCGGTAGCGGGTCCGGATCTCTGGACGGAAGCCCAGCTGGTTCC<br>CCTACGTCAACCGAAGAAGGAACCTCCGAATCCGCCACCCCCGAATCCGGTCCAGGAACCAGCACCGAACCTTCCGAGGGCAGCGCCCCGGGATC<br>GCCTGCGGGGAGCCCGACTTCCACTGAGGAAGGCACATCCACTGAGCCGTCCGAGGGATCAGCCCCCGGTACCAGCACTGAACCGAGCGAAGGAT<br>CGGCCCCCGGCACCTCCGAGTCTGCCACTCCGGAGTCCGGACCGGGGTCCGAGCCTGCAACTTCCGGGTCCGAAACTCCAGGATCGGAGCCTGCC<br>ACTAGCGGCAGCGAAACCCCGGGATCACCGGCCGGCTCCCCCACCTCGACCGAGGAAGGGACCTCCGAGAGCGCGACCCCTGAGTCGGGGCCCGG<br>AACCTCCACCGAACCCAGCGAAGGCTCGGCTCCCCTGGACGGCTCCGGCTCCGGGAGCGGAAGCGGATCAGGCTCCCTCGAGCATCATCACCACC |
| ISO-κ | *HEK293* (pROCK) | ATGGGATGGTCATGTATCATCCTTTTTCTAGTAGCAACTGCAACTGGAGTACATTCAGACATCCAGATGACCCAGTCTCCTTCACTCCTGTCTGC<br>ATCTGTGGGAGACAGAGTCACTCTCAACTGCAAAGCAAGTCAGAATATTAATAAGAACTTAGACTGGTATCAACAAAAGCTTGGAGAAGCGCCAA<br>AAGTCCTGATATATTATACAGACAATTTGCAAACGGGCTTCTCATCAAGGTTCAGTGGCAGTGGATCTGGTACAGATTACACACTCACCATCAGC<br>AGCCTGCAGCCTGAAGATGTTGCCACATATTACTGCTATCAGTATAACAGTGGGCCCACGTTTGGACCTGGGACCAAGCTGGAACTGAAACGGGC<br>TGATGCTGCACCAACTGTATCCATCTTCCCACCATCCAGTGAGCAGTTAACATCTGGAGGTGCCTCAGTCGTGTGCTTCTTGAACAACTTCTACC<br>CCAAAGACATCAATGTCAAGTGGAAGATTGATGGCAGTGAACGACAAAATGGCGTCCTGAACAGTTGGACTGATCAGGACAGCAAAGACAGCACC<br>TACAGCATGAGCAGCACCCTCACGTTGACCAAGGACGAGTATGAACGACATAACAGCTATACCTGTGAGGCCACTCACAAGACATCAACTTCACC<br>CATTGTCAAGAGCTTCAACAGGAATGAGTGTGCTAGCGGTGGTGGTGGTGGTGACTATAAGGACGACGACGACAAGTGA |

**Table S5. Additional mutant coiled-coil domains screened with elastin midblocks.** The Q domain was derived from P through fixation of 16 sequence mutations separately predicted to increase T cell-dependent immunogenicity. Two dozen related mutant coiled-coil domains were also screened for: 1) increased immunogenicity relative to P, and 2) the ability to promote hydrogel formation. During evolution, hydrogel formation was evaluated using an elastin-like midblock E. Triblocks formed using this midblock are susceptible to phase separation at low temperatures (LCST), particularly if the endblock coils aggregate. Predicted immunogenic domains generally had increased hydrophobicity and an increased tendency to aggregate. Only domains with an LCST > 37 °C were considered for biological studies. Each elastin-like hydrogel forming protein triblock was cloned and expressed in pQE-80L / *E. coli*, and had the following sequence:

| Domain | Length | Amino Acid Sequence |
|---|---|---|
| **E** (Elastin) | 154 | $\{[(VPGAG)_2VPGEG(VPGAG)_2]_3LD\}_2$ |
| Hydrogel forming triblock | 315 | $MRGSH_6GSVD(GS)_6G\{$**Coil**$\}(GS)_6LD\{$**E**$\}(GS)_6G\{$**Coil**$\}(GS)_6LEH_6KLN$* |

| Coil Name | Coil ID | Amino Acid Sequence | # Mutations |
|---|---|---|---|
| COMPcc = "P" | 1 | APQMLRELQETNAALQDVRELLRQQVKEITFLKNTVMESDAS | - |
| 4M | 2 | APQMLRELMETNMALQDVRELLRMQVKEITFLKMTVMESDAS | 4 |
| AMQA | 3 | APQMLRELQETNAALQDVREALRQQMKEITFLKNQVMESDAA | 4 |
| AA1 | 4 | APQMLRELQETNAALQDVRALLRQQVAEITFLKNTVMESDAS | 2 |
| AA2 | 5 | APQMLRELQETNAALQDVRELLAQQVKEITFLKNTVMASDAS | 2 |
| 5M6AQ | 6 | APQMLRELMETNMALQDVRAALAMQMAEITFLKMQVMASDAA | 12 |
| 1M6AQ | 7 | APQMLRELQETNAALQDVRAALAQQMAEITFLKNQVMASDAA | 8 |
| 1M8AQ | 8 | APQMLAELQATNAALQDVRAALAQQMAEITFLKNQVMASDAA | 10 |
| 5M6AQ-Q38 | 9 | APQMLRELQETNMALQDVRAALAMQMAEITFLKMQVMASDAA | 11 |
| 5M6AQ-A42 | 10 | APQMLRELMETNAALQDVRAALAMQMAEITFLKMQVMASDAA | 11 |
| 5M6AQ-Q53 | 11 | APQMLRELMETNMALQDVRAALAQQMAEITFLKMQVMASDAA | 11 |
| 5M6AQ-N63 | 12 | APQMLRELMETNMALQDVRAALAMQMAEITFLKNQVMASDAA | 11 |
| AMQA-L50 | 13 | APQMLRELQETNAALQDVRELLRQQMKEITFLKNQVMESDAA | 3 |
| AMQA-V55 | 14 | APQMLRELQETNAALQDVREALRQQVKEITFLKNQVMESDAA | 3 |
| AMQA-T64 | 15 | APQMLRELQETNAALQDVREALRQQMKEITFLKNTVMESDAA | 3 |
| AMQA-S71 | 16 | APQMLRELQETNAALQDVREALRQQMKEITFLKNQVMESDAS | 3 |
| Q2 | 17 | APQMLRELMATNMALQDVRAALAMQMAEITALKMQVMASDAA | 14 |
| QE1 | 18 | APQMERELMATNMALADVRAALAMQMAEIAALKMQVMASDEA | 18 |
| QE2 | 19 | AEEMLRELMATNMALADVRAALAMQMAEIAALKMQVMASDAA | 18 |
| QE3 | 20 | AEEMERELMATNMALADVRAALAMQMAEIAALKMQVMASDEA | 20 |
| Q7 = "Q" | 21 | AEEMLRELMATNMALADVRALLAQQMAEIAALKMQVMASDAA | 16 |
| Q8 | 22 | APEMLRELMATNMALADVRALLAQQMAEIAALKMQVMASDAA | 15 |
| Q10 | 23 | AEEMLRELMATNMALADVRALLAMQMAEIAALKMQVMASDEA | 18 |
| Q11 | 24 | AEEMERELMATNMALADVRALLAMQMAEIAALKMQVMASDAA | 18 |

**Table S6. Alignment of additional mutant coiled-coil domains.** Lines between heptads indicate structurally important positions that were not subject to mutational analysis. The heptad numbering is taken from Gunesakar et al.[11]

| coil ID # | heptad | a | b | c | d | e | f | g | a | b | c | d | e | f | g | a | b | c | d | e | f | g | a | b | c | d | e | f | g | a | b | c | d | e | f | g | a | b | c | d | e | f | g | |
|---|---|---|---|---|---|---|---|---|---|---|---|---|---|---|---|---|---|---|---|---|---|---|---|---|---|---|---|---|---|---|---|---|---|---|---|---|---|---|---|---|---|---|---|---|
| | numbering | 30 | 31 | 32 | 33 | 34 | 35 | 36 | 37 | 38 | 39 | 40 | 41 | 42 | 43 | 44 | 45 | 46 | 47 | 48 | 49 | 50 | 51 | 52 | 53 | 54 | 55 | 56 | 57 | 58 | 59 | 60 | 61 | 62 | 63 | 64 | 65 | 66 | 67 | 68 | 69 | 70 | 71 | # mut |
| 1 | "P" COMPcc | A | P | Q | M | L | R | E | L | Q | E | T | N | A | A | L | Q | D | V | R | E | L | L | R | Q | Q | V | K | E | I | T | F | L | K | N | T | V | M | E | S | D | A | S | - |
| 2 | 4M | A | P | Q | M | L | R | E | L | M | E | T | N | M | A | L | Q | D | V | R | E | L | L | R | M | Q | V | K | E | I | T | F | L | K | M | T | V | M | E | S | D | A | S | 4 |
| 3 | AMQA | A | P | Q | M | L | R | E | L | Q | E | T | N | A | A | L | Q | D | V | R | E | A | L | R | Q | Q | M | K | E | I | T | F | L | K | N | Q | V | M | E | S | D | A | A | 4 |
| 4 | AA1 | A | P | Q | M | L | R | E | L | Q | E | T | N | A | A | L | Q | D | V | R | A | L | L | R | Q | Q | V | A | E | I | T | F | L | K | N | T | V | M | E | S | D | A | S | 2 |
| 5 | AA2 | A | P | Q | M | L | R | E | L | Q | E | T | N | A | A | L | Q | D | V | R | E | L | L | A | Q | Q | V | K | E | I | T | F | L | K | N | T | V | M | A | S | D | A | S | 2 |
| 6 | 5M6AQ | A | P | Q | M | L | R | E | L | M | E | T | N | M | A | L | Q | D | V | R | A | A | L | A | M | Q | M | A | E | I | T | F | L | K | M | Q | V | M | A | S | D | A | A | 12 |
| 7 | 1M6AQ | A | P | Q | M | L | R | E | L | Q | E | T | N | A | A | L | Q | D | V | R | A | A | L | A | Q | Q | M | A | E | I | T | F | L | K | N | Q | V | M | A | S | D | A | A | 8 |
| 8 | 1M8AQ | A | P | Q | M | L | A | E | L | Q | A | T | N | A | A | L | Q | D | V | R | A | A | L | A | Q | Q | M | A | E | I | T | F | L | K | N | Q | V | M | A | S | D | A | A | 10 |
| 9 | 5M6AQ-Q38 | A | P | Q | M | L | R | E | L | Q | E | T | N | M | A | L | Q | D | V | R | A | A | L | A | M | Q | M | A | E | I | T | F | L | K | M | Q | V | M | A | S | D | A | A | 11 |
| 10 | 5M6AQ-A42 | A | P | Q | M | L | R | E | L | M | E | T | N | A | A | L | Q | D | V | R | A | A | L | A | M | Q | M | A | E | I | T | F | L | K | M | Q | V | M | A | S | D | A | A | 11 |
| 11 | 5M6AQ-Q53 | A | P | Q | M | L | R | E | L | M | E | T | N | M | A | L | Q | D | V | R | A | A | L | A | Q | Q | M | A | E | I | T | F | L | K | M | Q | V | M | A | S | D | A | A | 11 |
| 12 | 5M6AQ-N63 | A | P | Q | M | L | R | E | L | M | E | T | N | M | A | L | Q | D | V | R | A | A | L | A | M | Q | M | A | E | I | T | F | L | K | N | Q | V | M | A | S | D | A | A | 11 |
| 13 | AMQA-L50 | A | P | Q | M | L | R | E | L | Q | E | T | N | A | A | L | Q | D | V | R | E | L | L | R | Q | Q | M | K | E | I | T | F | L | K | N | Q | V | M | E | S | D | A | A | 3 |
| 14 | AMQA-V55 | A | P | Q | M | L | R | E | L | Q | E | T | N | A | A | L | Q | D | V | R | E | A | L | R | Q | Q | V | K | E | I | T | F | L | K | N | Q | V | M | E | S | D | A | A | 3 |
| 15 | AMQA-T64 | A | P | Q | M | L | R | E | L | Q | E | T | N | A | A | L | Q | D | V | R | E | A | L | R | Q | Q | M | K | E | I | T | F | L | K | N | T | V | M | E | S | D | A | A | 3 |
| 16 | AMQA-S71 | A | P | Q | M | L | R | E | L | Q | E | T | N | A | A | L | Q | D | V | R | E | A | L | R | Q | Q | M | K | E | I | T | F | L | K | N | Q | V | M | E | S | D | A | S | 3 |
| 17 | Q2 | A | P | Q | M | L | R | E | L | M | A | T | N | M | A | L | Q | D | V | R | A | A | L | A | M | Q | M | A | E | I | T | A | L | K | M | Q | V | M | A | S | D | A | A | 14 |
| 18 | QE1 | A | P | Q | M | E | R | E | L | M | A | T | N | M | A | L | A | D | V | R | A | A | L | A | M | Q | M | A | E | I | A | A | L | K | M | Q | V | M | A | S | D | E | A | 18 |
| 19 | QE2 | A | E | E | M | L | R | E | L | M | A | T | N | M | A | L | A | D | V | R | A | A | L | A | M | Q | M | A | E | I | A | A | L | K | M | Q | V | M | A | S | D | A | A | 18 |
| 20 | QE3 | A | E | E | M | E | R | E | L | M | A | T | N | M | A | L | A | D | V | R | A | A | L | A | M | Q | M | A | E | I | A | A | L | K | M | Q | V | M | A | S | D | E | A | 20 |
| 21 | "Q" Q7 | A | E | E | M | L | R | E | L | M | A | T | N | M | A | L | A | D | V | R | A | L | L | A | Q | Q | M | A | E | I | A | A | L | K | M | Q | V | M | A | S | D | A | A | 16 |
| 22 | Q8 | A | P | E | M | L | R | E | L | M | A | T | N | M | A | L | A | D | V | R | A | L | L | A | Q | Q | M | A | E | I | A | A | L | K | M | Q | V | M | A | S | D | A | A | 15 |
| 23 | Q10 | A | E | E | M | L | R | E | L | M | A | T | N | M | A | L | A | D | V | R | A | L | L | A | M | Q | M | A | E | I | A | A | L | K | M | Q | V | M | A | S | D | E | A | 18 |
| 24 | Q11 | A | E | E | M | E | R | E | L | M | A | T | N | M | A | L | A | D | V | R | A | L | L | A | M | Q | M | A | E | I | A | A | L | K | M | Q | V | M | A | S | D | A | A | 18 |

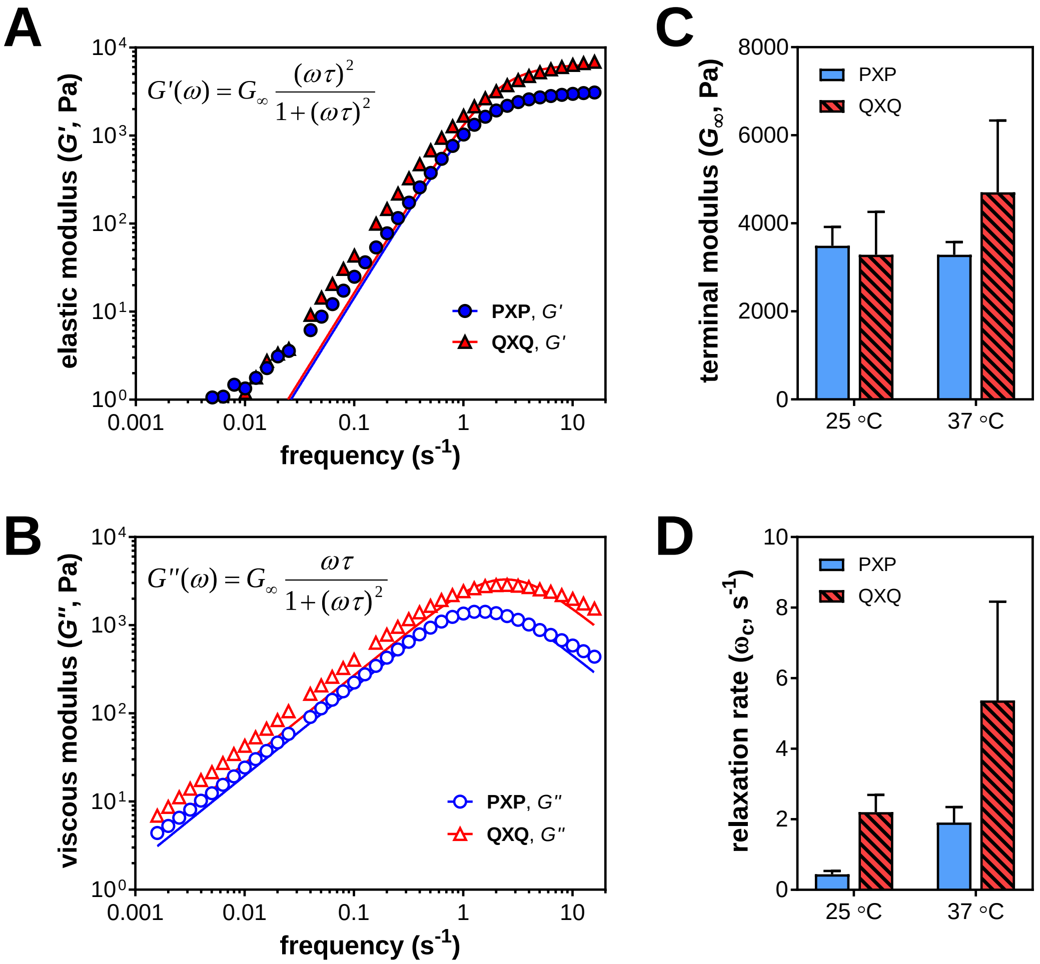

**Figure S1. Summary of rheological properties of PXP and QXQ hydrogels.** Hydrogels were prepared by swelling lyophilized mesh proteins in 100 mM phosphate buffer, pH 7.4 at a fixed concentration of 10% (*w*/*v*). (**A**) Frequency response of the elastic modulus (*G*') at 37 °C. Frequency sweeps were performed at a fixed strain amplitude of 1% between 0.01 and 100 rad $s^{-1}$. The solid lines represent best fits to **eq. S1**. (**B**). Frequency response of the viscous modulus (*G*'') at 37 °C. The solid lines represent best fits to **eq. S2**. (**C**) The terminal moduli ($G_\infty$) of the gels at 25 °C and 37 °C, extracted from best fits. (**D**) The relaxation rate $\omega_c$ for the gels at 25 °C and 37 °C, extracted from best fits. This rate represents the crossover frequency between *G*' and *G*''. Points in each graph represent μ ± σ for 3 independent gel preparations per temperature.

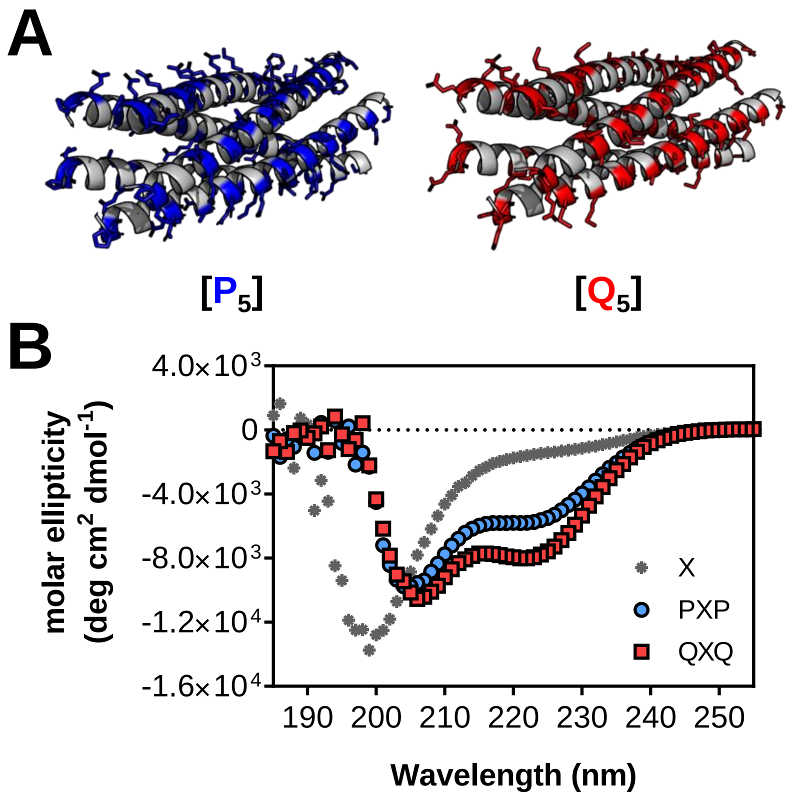

**Figure S2. Circular dichroism spectroscopy of dilute solutions of artificial proteins.** (**A**) AF2-predicted structural model of the $P_5$ and $Q_5$ pentameric coiled-coil bundles (see also PDB 1VDF). The predicted structures align well with a low RMSD of 0.351 $Å^2$. The pentamerized coils are hypothesized to serve as network junctions in the self-assembled hydrogels. Surface exposed sidechains are highlighted. (**B**) Circular dichroism spectra of dilute hydrogel-forming proteins, together with the non-gel forming X domain. The protein samples were prepared at 30 µM (ca. 1 mg $mL^{-1}$) in 100 mM phosphate buffer, 0.2 µm-filtered, and scanned at room temperature between 185 nm and 300 nm in 1 nm wavelength step increments with a spectral averaging time of 5 seconds per step. Shown are representative ellipticity traces for two independent sample preparations per protein.

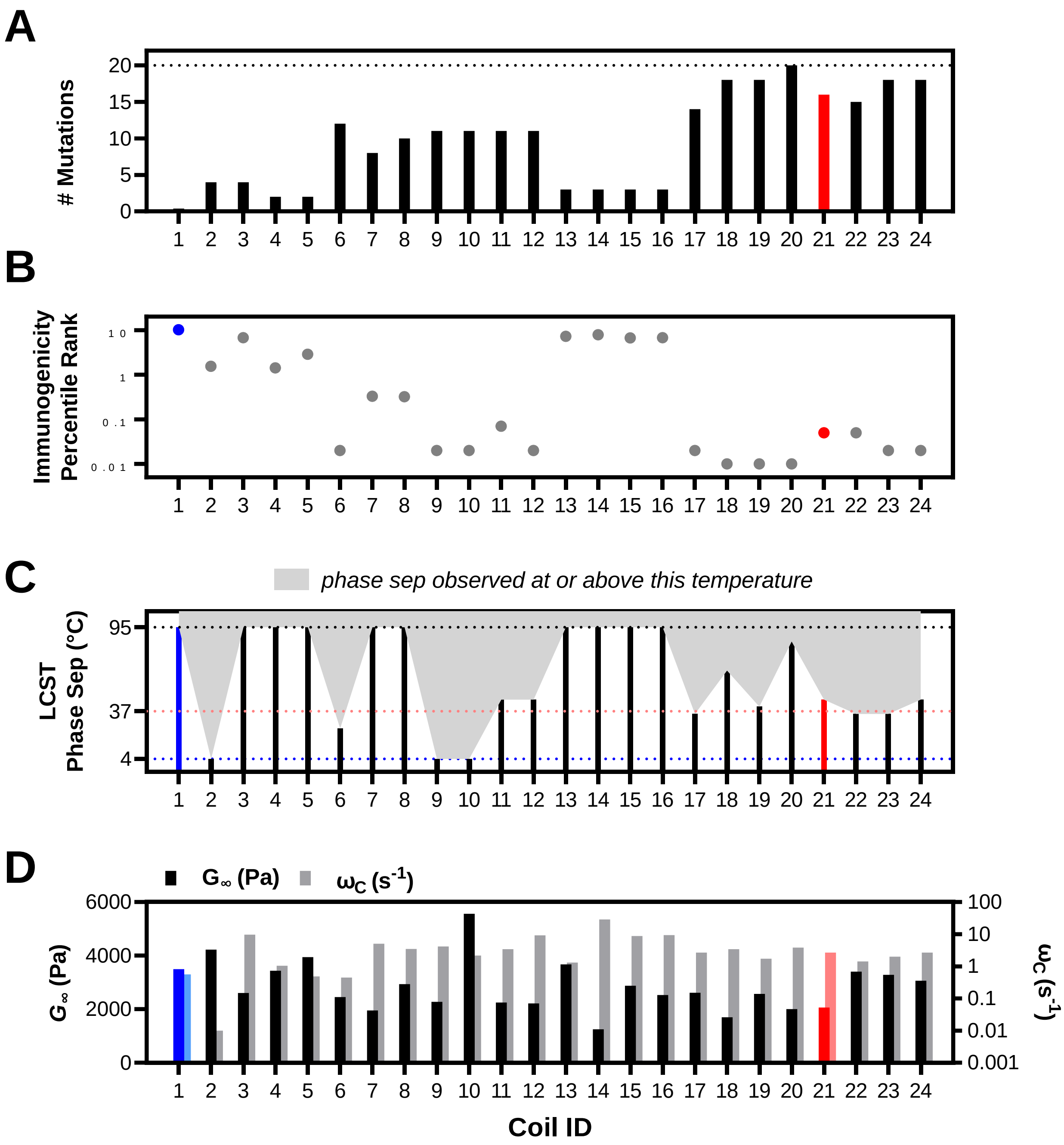

**Figure S3. Properties of additional mutant coiled-coil domains**. The coiled-coil forming domains in **Table S5** were screened for their potential immunogenicity, solubility, and ability to self-assemble into a viscoelastic hydrogel network. (**A**) Mutational distance from the starting P domain for coil sequences shown in **Table S5**. (**B**) Immunogenicity of each coil as predicted by the NIH Immune Epitope Database (IEDB). The predicted immunogenicity increases with decreasing percentile rank. Shown are the average consensus percentile ranks for the top five peptides and their corresponding peptide binding registers within the MHC II allele H2 I-$A^d$. For context, ovalbumin (OVA) protein presents several model complexes within this allele, and the average percentile rank for the top five OVA peptides was 0.09. (**C**) Each coiled-coil was screened for its ability to drive the formation of ordered hydrogels when used as symmetric endblocks flanking a flexible midblock derived from elastin. Each hydrogel-forming triblock had the following sequence: $\texttt{MRGSH}_6\texttt{GSVD(GS)}_6\texttt{G[}\textbf{Coil}\texttt{](GS)}_6\texttt{LD[}\textbf{E}\texttt{](GS)}_6\texttt{G[}\textbf{Coil}\texttt{](GS)}_6\texttt{LEH}_6\texttt{KLN*}$ (where **E** represents the elastin sequence from **Table S5**). In general, block hydrophobicity was found to increase with predicted immunogenicity. Hydrophobic blocks tend to be more disordered, and can more easily phase separate at a lower critical solution temperature (LCST). The final selected sequence, Coil #21 (a.k.a. "Q") was found to have an LCST > 37 °C, indicating that gels formed from this domain would be homogenously swollen at biological temperatures. Note that although coil-elastin-coil (CEC) triblocks showed such phase separation behavior, the XTEN-based CXC triblocks did not show this behavior. This lack of thermosensitive phase behavior for the XTEN gels was an additional motivating factor in selecting it for biological studies. The temperature range of the assay was between 4 and 95 °C, and gels with LCST outside of this range are plotted on the dotted lines. For Coil #6 (5M6AQ), the gel had an LCST less than 25 °C. (**D**) Linear rheology of the CEC triblocks at room temperature, similar to Figure S2. The terminal modulus $G_\infty$ and characteristic network relaxation time τ were extracted from fits to *G'* (elastic modulus) and *G"* (viscous modulus) as a function of frequency using eq. **S1** and **S2**. The relaxation time is related to the characteristic relaxation rate $\omega_c = 1 / \tau$. Due to the large amount of protein powder required for each rheological test, as well as the large number of CEC-type proteins tested, each gel in panel C and D was screened for phase separation and viscoelasticity only once ($N = 1$ experimental replicate). The resultant property dataset was used primarily to aid in selection of Coil #21 as the final "non-self" coiled-coil domain.

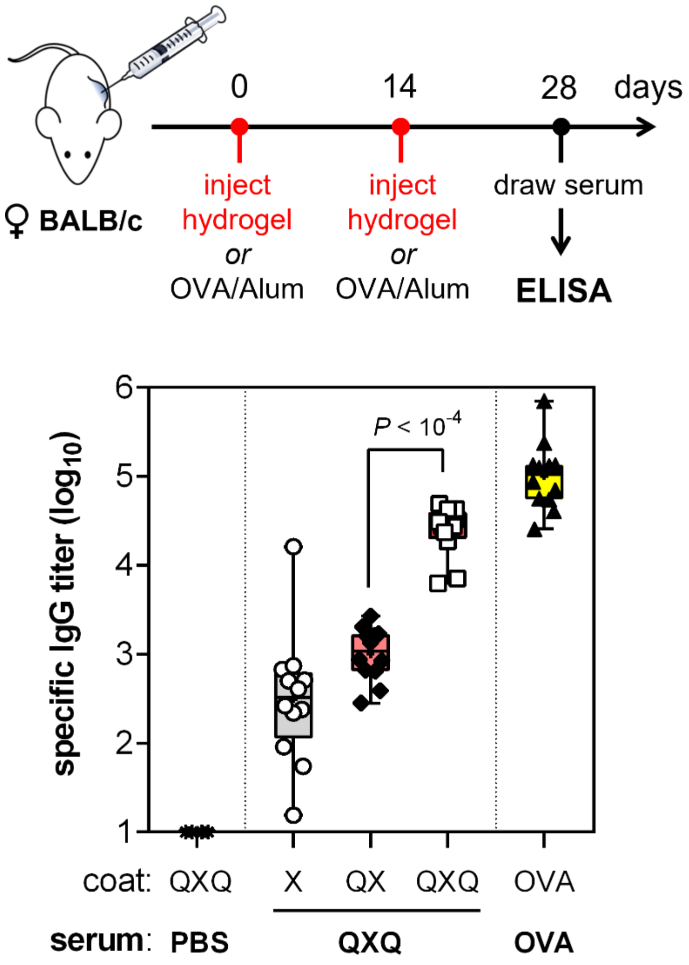

**Figure S4. QXQ hydrogels induce similar antibody titers as Ovalbumin.** Mice were challenged twice with subcutaneous QXQ hydrogel implantation (SC, 5 mg, 10% *w*/*v* in 50 µL) or by OVA/Alum (IP, 50 µg) on d0 or d14. On d28, serum was collected and assayed for specific IgG titers by antigen-specific ELISA. Mice from QXQ-challenged groups exhibited low, intermediate and high levels of antibody titers against X, QX, and QXQ, respectively. The QXQ response was similar in magnitude to the response observed against OVA/Alum, which is commonly considered to be a strong antigen. Shown are all serum results obtained from 3 independent groups of 4 mice/group ($n$ = 12 mice total).

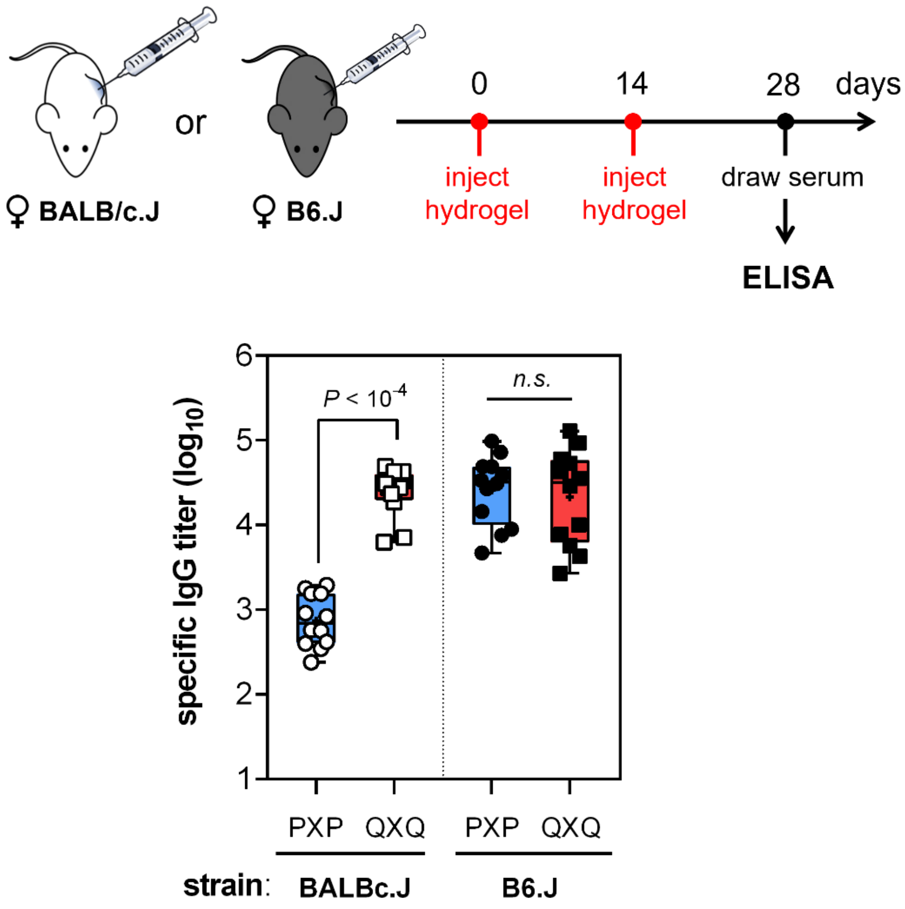

**Figure S5. Strain-specific response to QXQ hydrogels.** Female BALB/c.J or B6.J mice (8 – 10 wks old) were challenged twice with subcutaneous hydrogel implantation (PXP or QXQ) on d0 and d14. Serum was withdrawn on d28 and assayed for specific IgG titers by antigen-specific ELISA. Shown are all serum results obtained from 3 independent groups of 4 mice/group ($n$ = 12 mice total). Whereas BALBc.J mice showed a strong and reproducible titer difference between PXP and QXQ, B6.J mice mounted similarly high titers against both PXP and QXQ. This strain-dependent reactivity difference is consistent with the design criterion for Q, which was evolved to have high affinity for MHC II I-$A^d$. This MHC II haplotype is not found in the B6 mouse. Data for the BALBc.J strain is the same as in **Figure 3** of the main text.

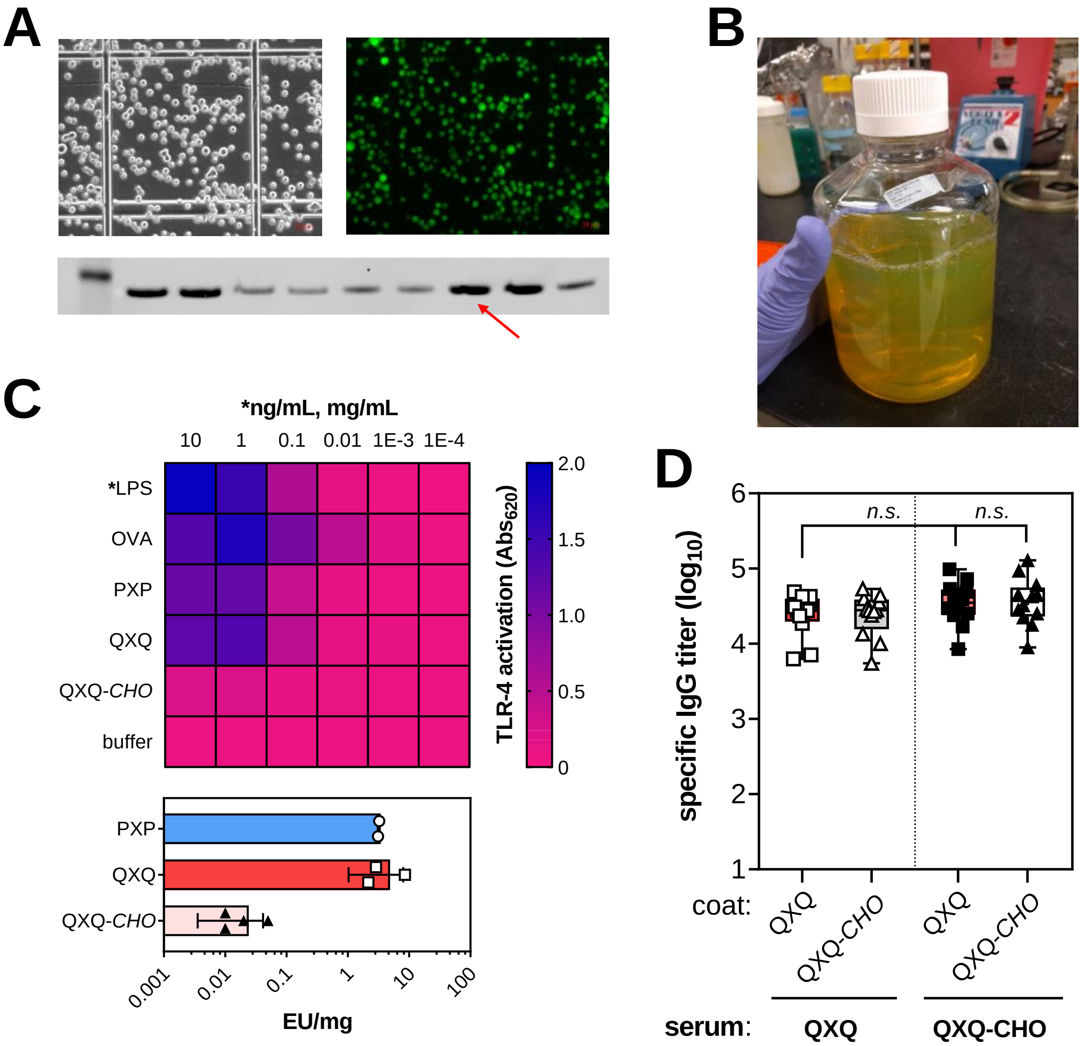

**Figure S6. Removal of endotoxin does not blunt IgG response against QXQ.** (**A**) Fluorescent micrographs of ExpiCHO cells stably transfected with pD2537 plasmid carrying an QXQ-IRES-GFP cassette (see *Materials and Methods*) for details. The anti-His western blot against QXQ highlights select high-expressing clones (e.g. 260 mg $L^{-1}$ expression level from one clone, red arrow) (**B**) Clarified supernatant from a 1 L expression of QXQ in ExpiCHO cells carrying QXQ-IRES-GFP. (**C**) Endotoxin levels from *CHO*-derived QXQ were <0.1 EU/mg as determined by a TLR-4 HEK-Blue activation assay. (**D**) Removal of endotoxin did not blunt the observed high titer response against plated QXQ antigen. Female BALB/c.J (8 – 10 wks old) were challenged twice with subcutaneous hydrogel implantation (QXQ or QXQ-*CHO*) on d0 and d14. Serum was withdrawn on d28 and assayed for specific IgG titers by antigen-specific ELISA. Shown are all serum results obtained from 3 independent groups of 4 mice/group ($n$ = 12 mice total).

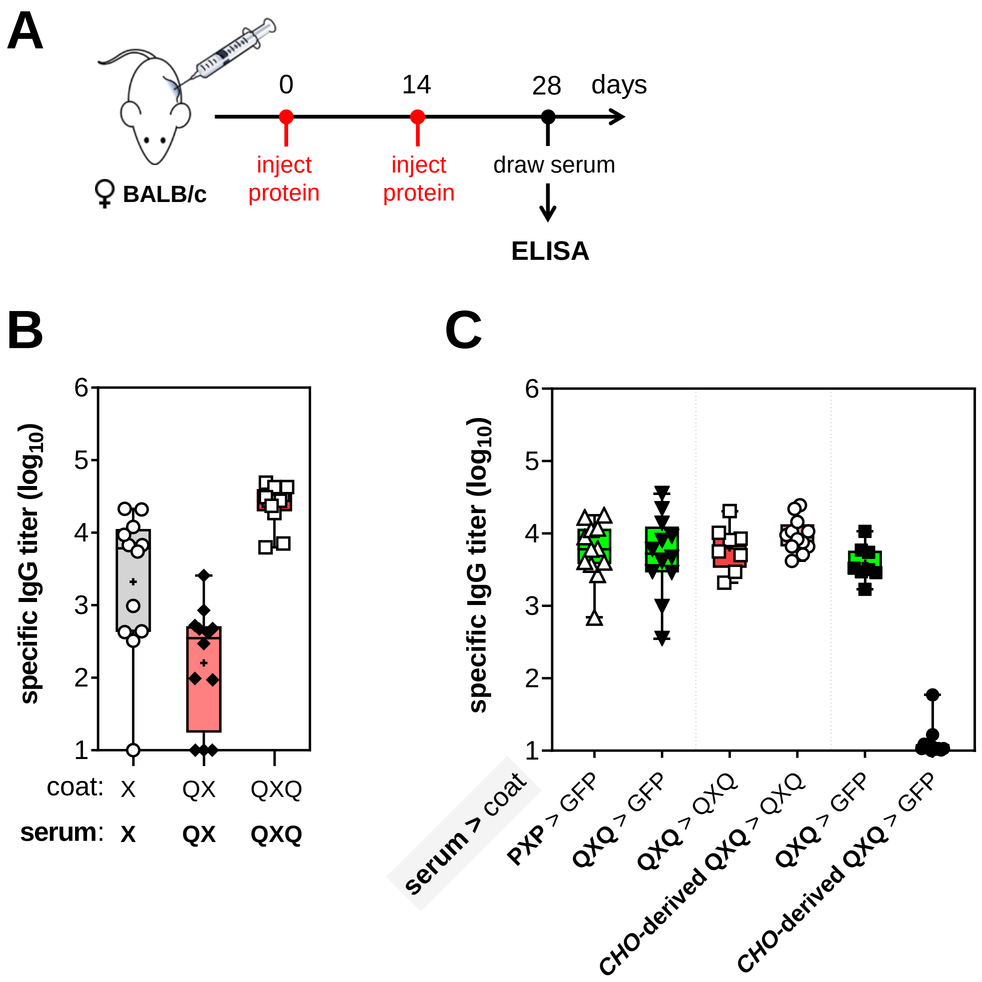

**Figure S7. Analysis of epitope specificity and expression species reactivity.** (**A**) Female BALB/c.J (8 – 10 wks old) were challenged twice with protein injection (X, QX, QXQ or PXP) on d0 (prime) and d14 (boost). Serum was withdrawn on d28 and assayed for specific IgG titers by antigen-specific ELISA. (**B**) Injection of *E. coli*-derived X or QX (the proteins each carry 6xHis at both N- and C-termini) induced intermediate IgG titers against the same coat protein. (**C**) Sera from both PXP- and QXQ-challenged mice cross-reacted with *E. coli*-derived 6xHis-tagged GFP coat protein, whereas the cross-reactivity against *E. coli*-derived GFP coat protein was undetectable when challenge sera from *CHO*-derived QXQ was plated. Shown are all serum results obtained from 2 - 3 independent groups of 4 mice/group (*n* = 8 - 12 mice total).

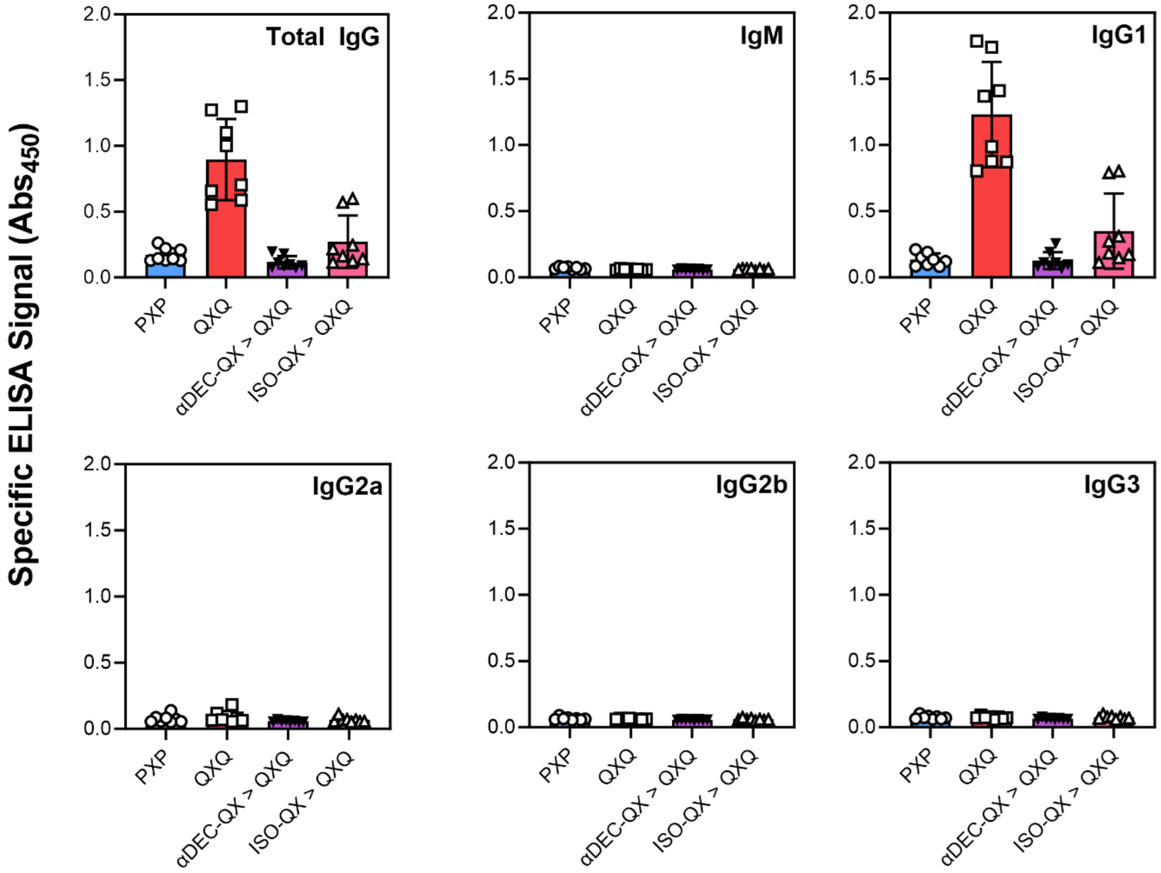

**Figure S8. Antibody isotyping of recalled sera.** Mice were prophylactically treated with either αDEC-QX, ISO-QX, or left untreated before being challenged with hydrogel injection (either QXQ or PXP), as in Fig. 4. Serum was withdrawn on d28 and assayed for coat-specific antibody titers by antigen-specific ELISA. Individual isotypes were quantified with heavy-chain anti-mouse antibodies (Abcam). Shown are all serum results obtained from two independent groups of 4 mice/group, representative of five such groups ($n$ = 20 mice total per group). All sera were assayed at a dilution of $1{:}10^4$.
